# Heritable morphology-environment correlations among lake populations of threespine stickleback

**DOI:** 10.64898/2026.08.20.745995

**Authors:** Alex Yeung, Ben A. Flanagan, Heather Alexander, Emma S. Choi, John Berini, Abigail G. Albright, Caroline Szajada, Nataly Vargas, Julian Flanagan, Edith Reyes Contreras, Peaches Cooper, Mehreen Shahid, Penelope R. Steffen, Fahad Gilani, Ana Santacruz, Viola Watts, Emma Polard, Kaligua Rochon, Emma Redfield, Jessica Hite, Amanda K. Hund, Daniel I. Bolnick

## Abstract

Phenotypic differences among populations can arise through heritable genetic divergence, phenotypic plasticity, or both, making it difficult to determine whether trait-environment correlations observed in nature reflect adaptive evolution. Within threespine stickleback (*Gasterosteus aculeatus*) studies, numerous document morphological differences among allopatric-, parapatric-, and even sympatric populations. These phenotypic differences among populations are often correlated with diet and lake habitat (e.g., lake size), suggesting an adaptive value to the population differences. However, many studies of ecomorphological divergence in stickleback use wild-caught stickleback, which may differ due to evolution or plasticity. Although common garden experiments have confirmed that population differences can be heritable, such experiments typically entail small numbers of populations. Consequently, we still do not know to what extent well-known trait-environment correlations in stickleback are a result of evolution. To address this gap, we reared stickleback embryos from 27 lake populations on Vancouver Island, in a laboratory environment. Morphological differences among populations persist in common-garden fish, confirming a large role for divergent evolution. These heritable differences were associated with environmental variation among lakes, implying an adaptive value. However, some well-known trait-environment relationships in stickleback did not persist in common-garden fish and may be primarily plastic.

## Introduction

Isolated populations of a given species can be phenotypically distinct (Forsman, 2015), either because populations have evolved in response to different environments or because different environments induce plastic differences among populations (West-Eberhard, 2003; Wimberger, 1991). A central goal of evolutionary biology is therefore to determine the relative contributions of heritable genetic divergence and phenotypic plasticity to morphological differences among populations. Common garden experiments provide a powerful approach for distinguishing these alternatives. In a common garden experiment, individuals of different populations are reared under standardized environmental conditions, minimizing environmental differences among populations (Ballentine and Greenberg, 2010; Euclide et al., 2022; Campbell et al., 2022; Martínez-Sancho et al., 2025). Differences among populations raised under similar environmental conditions provide evidence that phenotypic divergence has a heritable basis. Thus, common garden experiments can determine whether population differences observed in the wild persist when populations experience similar environments.

Another central goal of evolutionary biology is to determine whether phenotypic differences among populations are adaptive. If natural selection drives repeated evolution of particular traits in a given environment, then that trait will be correlated with the relevant environmental variation. Therefore, trait-environment correlations are widely invoked as evidence of adaptive divergence. However, phenotypic plasticity can also produce trait-environment correlations. Do among-population trait-environment correlations persist in a common garden setting? If so, this would be evidence that the correlations are heritable, and likely represent adaptive evolution. If not, the correlations are likely plastic.

Common garden rearing has long been used to test for heritable among-population differences, especially in plants and invertebrates that are readily bred and reared in large numbers in a greenhouse or laboratory setting. Baughman et al. (2019) reviewed 121 plant taxa, 170 studies, and 327 experiments. They found that most experiments reported differences among populations and reported trait-environment correlations. Small animals such as flies (Misra and Reeve, 1964; 12 populations) and mice (Phifer-Rixey et al, 2018; 5 populations) have been reared in common gardens to study trait environment correlations, but to a generally smaller scale. Differences among populations and trait-environment correlations were also found in these animals. Unfortunately, studies of larger animals, especially vertebrates, rarely do large many-population common garden experiments. Therefore, for many organisms we do not know whether trait-environment correlations actually represent heritable adaptations. For example, the very intensively studied evolutionary model organism, threespine stickleback (*Gasterosteus aculeatus*), have been used in many common garden experiments. But, these studies generally use only a few populations (Supplementary Table S1), which limits our ability to test trait-environment correlations rigorously.

Here, we use a large common garden experiment to assess heritable morphological variation and trait-environment correlations stickleback. Stickleback evolved morphological adaptations to contrasting environments following their expansion from marine into freshwater habitats after Pleistocene deglaciation (∼11,000 years). For example, freshwater stickleback populations are smaller, have fewer armor plates, and have fewer, shorter gill rakers when compared to marine populations (Reimchen, 1983; Bell and Foster, 1994; McKinnon and Rundle, 2002; Colosimo et al., 2005). Stickleback also evolved divergent adaptations to distinct lake habitats. Sympatric benthic and limnetic species pairs are adapted to shallow littoral and pelagic habitats, respectively (Schluter and McPhail, 1992). Limnetic stickleback are heritably smaller and slimmer, have more armor plates, have smaller mouths, and have longer, more numerous gill rakers compared to benthic stickleback (Schluter and McPhail, 1992; MacColl, 2009). However, most lakes contain single populations of stickleback, not species pairs. These ‘solitary’ populations vary in morphology, spanning a gradient from relatively benthic-like to relatively limnetic-like morphology (Schluter, 1995; Hatfield and Schluter, 1999). This among-population morphological diversity is often correlated with lake habitats (e.g., lake surface area), suggesting an adaptive value. However, trait-environment correlations across populations have generally been studied using wild-caught stickleback, thereby conflating the roles of plasticity and heritability.

Stickleback studies demonstrate a strong genetic basis for between-population differences in body shape (Reid and Peichel, 2010), gill raker number (Hagen, 1973; McPhail, 1992), gill raker length (Lavin & McPhail, 1987; Schluter, 1996; Hatfield, 1997), and armor plate number (Colosimo et al., 2005). However, other studies of stickleback reveal phenotypic plasticity in response to diet (Day et al., 1994; Svanbäck & Schluter, 2012; Lindsey, 1962; Day et al., 1994; Hairston et al., 1982; Rick et al., 2012; Willacker et al., 2010), predation (Ab Ghani et al., 2016; Miller et al., 2015; Howland et al., 2004; Voje et al., 2013), parasite regime (Stutz et al., 2015), and light regime (Veen et al., 2017). Consequently, trait-environment correlations measured on wild-caught stickleback may reflect either adaptive evolution, plasticity, or both.

Here, we present a large-scale common garden study of morphological divergence of 27 lake populations of stickleback. First we ask, are there inherited among-population differences in morphology that persist in the lab-reared fish? Second we ask, are the heritable population differences correlated with environmental features of their native lakes, implying adaptive evolution? We expect all morphological traits to have differences among populations and for traits to be correlated with environmental features along the major benthic-limnetic axis.

In addition to these core questions, we also evaluate the magnitude of sexual dimorphism in stickleback ecomorphology, and test whether this differs between populations in a common-garden setting. Stickleback exhibit strong dimorphism in body shape (Kitano et al., 2007; Aguirre et al., 2008), gill raker morphology (Bolnick and Lau 2008), diet (Bolnick et al., 2014), and many other traits (Reimchen and Nosil, 2004; Aguirre and Akinpelu, 2010; Leinonen 2011; Mcgee and Wainwright, 2013). This sexual dimorphism is commonly interpreted as an adaptation to specific sex reproductive roles driven by mate choice and competition (Greenwood & Adams, 1987; Andersson & Iwasa, 1996; Olsson et al., 2002; Hamon & Foote, 2005; Roy et al., 2007; De Lisle & Rowe, 2014; Keagy et al., 2016; Head et al., 2013). Ecological partitioning between the sexes may also contribute to dimorphism (Shaffer et al., 2001; Kitano et al., 2007; Bolnick and Lau 2008). The degree of sexual dimorphism has been found to differ between populations and habitats (Spoljaric & Reimchen, 2008; Bolnick and Lau, 2008; Leinonen et al., 2011). However, studies of sexual dimorphism in stickleback are rarely conducted in common garden settings, so it remains unclear to what extent among-population variation in the magnitude of dimorphism is heritable.

## Materials and Methods

### Fish collection

In late May and early June 2024, we used minnow traps and seine nets to capture stickleback from 27 lakes on Vancouver Island, Canada (Supplementary Table S2), retaining gravid females and mature males. The 27 lakes were chosen to represent several geographic regions of the Island (west coast, east coast, and northern), and a range of lake sizes and features within each region (Supplementary Figure S1). Collection was permitted by the Ministry of Forest, Lands, and Natural Resources (permit NA24-89569), the Huu-ay-aht First Nations (permit 2024-042), and the University of Connecticut IACUC (protocol A21-025).

### Within-population crosses and husbandry

We generated 5-10 full-sib families of embryos from each of the 27 lakes. Wild-caught gravid females and adult males were euthanized in MS-222. We gently squeezed females to deposit eggs into a petri dish. We quickly added sperm from testes, dissected from males and macerated in a drop of water (Spence-Jones, 2024). Fertilized eggs were rinsed after a minute, then transported to the Bamfield Marine Sciences Center (BMSC; AUP protocol #RS-22-09) in 50 mL Falcon tubes.

Each full-sib family of embryos were placed in a large Petri dish and immersed in water with dilute methylene blue (changed daily). Once the embryos hatched, we moved larvae into flow-through, aerated 3L Z-hab tanks with automated 20% daily dechlorinated municipal water. After swim-up and yolk sac depletion, stickleback larvae were fed live dechorionated artemia larvae twice daily.

Once individual stickleback were approximately 1.5 cm long, we transferred them to flow-through 38L glass aquariums with artificial plants for enrichment, a biological sponge filter, and automated 10% water changes six times daily. Once stickleback reached about 2 cm long we transitioned them to a ground mix of dry food consisting of freeze-dried mysis shrimp, freeze-dried blood worms, Gemma Micro (800µm), and Golden Pearls (500-800µm) supplemented with artemia. The water temperature in the aquariums fluctuated seasonally from 15°C (summer) down to 7.5°C (winter). The light regime was maintained at 16:8 hours (light:dark).

### Dissections, photographs, and measurements

In March 2025, we euthanized all stickleback in 50 mg/L neutral buffered MS-222. Sample sizes for each lake are provided in Supplementary Table S2. The carcasses were weighed to the nearest 0.01g. We photographed the right side of each fish against a standard background with a ruler for scale, with consistent light and camera settings. Using ImageJ Version 1.54p 17, we subsequently used the whole-body photographs to measure standard length, head length, snout length, eye diameter, caudal peduncle depth, and body depth (Stuart et al., 2017). We recorded sex by inspecting gonads after dissection, then fixed each in 10% neutral buffered formalin for a few weeks. The fixed fish were subsequently rinsed, stained in alizarin red for two days, then stored in 70% isopropyl alcohol.

We used calipers to measure the gape width of stained specimens. Armor plates were counted on the left side. We took a photograph of the stained neurocranium, and later digitally measured lengths of the neurocranial elements of the opercular 4-bar lever (Westneat, 1990; Wainwright et al., 2005; Thompson et al., 2017). These lengths were used to calculate opercular 4-bar kinematic transmission (KT), a measure of jaw opening performance (as described in Thompson et al, 2017). Removing the operculum, we counted gill rakers on the first left branchial arch. Then we removed and photographed the arch with a micrometer under a dissecting microscope, to digitally measure length of the two longest gill rakers and raker density (average spacing between five consecutive rakers). The pectoral fin was cut off, splayed, and photographed to digitally measure width and length. These measurements were used to calculate the pectoral fin shape (log[length/width]). This shape is an approximation of the functionally important aspect ratio (Wainright et al, 2002). In total, we measured 14 morphological traits on all 27 lake populations.

### Limnological and biotic variables

Biotic and abiotic characteristics of the 27 lakes were quantified, as previously described by Berini et al. (2026), Srinivas et al. (2026) and Choi et al (2026). To briefly summarize, we used iMapBC, an interactive mapping program hosted by the government of British Columbia, Canada, to extract bathymetry data for each lake including perimeter (m), maximum depth (m), mean depth (m), surface area (m^2^), and elevation (m). We employed an EXO2 multi-probe sonde (YSI Incorporated, Ohio, USA) to measure temperature-depth profiles at one-meter increments to one-meter above the hypolimnion. Profiles were collected at each lake’s deepest point to a maximum depth of 20 meters. The YSI also generated depth profiles for pH, dissolved oxygen (mg L-1; also computed as % saturation), and chlorophyll a (a stand-in for phytoplankton biomass, measured in relative fluorescence units (RFUs)).

Zooplankton tows (late May through June 2023) provided samples of copepod abundance and community composition in the wild. At each lake, we collected two zooplankton samples from different locations, with each sample consisting of three bottom-to-surface tows using a Wisconsin net. Samples were preserved and later used to estimate aerial densities of copepods and other zooplankton taxa, as well as overall zooplankton community diversity (Srinivas et al, 2026). Choi et al., (2026) obtained estimates of infection prevalence by a major parasite (*Schistocephalus solidus*) in the wild. This parasite is transmitted via limnetic prey (cyclopoid copepods), so its prevalence tends to be higher in stickleback individuals and populations with a more limnetic diet (Stutz et al., 2014; Weber et al., 2022). We also included the average severity of peritoneal fibrosis in each population in the wild (from Choi et al. (2026), which contributes to resistance to *S. solidus* (Weber et al. 2022; Bolnick et al., 2026). Sample sizes used for fibrosis and infection studies are provided in Supplementary Table S2.

### Analyses

Prior to statistical analyses, we size-standardized the linear univariate measures of morphology, retaining residuals of log-transformed traits regressed on log standard length, except for standard length, armor plate number, gill raker number, and fin aspect ratio. All analyses and visualizations were done in R version 2026.01.0+392 (R Core Development Team, 2025). The raw data, R scripts, and metadata are provided in a data repository on Figshare (doi: 10.6084/m9.figshare.c.8665041).

### Among-population differences in morphology

To test for among-population differences in multivariate morphology among lab-raised fish, we applied a Multivariate Analysis of Variance (MANOVA) to all 14 morphological variables. A linear discriminant analysis (LDA) allowed us to identify combinations of morphological traits that best discriminate among populations. A confusion analysis allowed us to understand how the different populations are sorted in the LDA. Additionally, we used an Analysis of Variance (ANOVAs) to test for among-population differences in each univariate trait. For this analysis we determined significance based on a Sequential Bonferroni corrected *p*-value with a default alpha of 0.05. To test for population differences in allometry, we used linear models in which each univariate trait was regressed on log standard length, with effects of population and a length x population interaction. Statistical significance was evaluated using Type III Sums of Squares. We retained the slope of each trait-length relationship for each lake. To test whether population divergence entails integrated combinations of traits, we calculated population means for every trait. We then calculated Pearson correlations between population means, for each pairwise combination of traits.

### Sexual dimorphism

We tested whether sexual dimorphism in morphology persists in lab-raised stickleback. We used a linear model to evaluate each trait’s dependence on length, sex, population, and their interaction (with Type III Sums of Squares). The main effect of sex tests for a consistent direction of dimorphism that holds across populations. The population x sex interaction tests whether the magnitude or direction of dimorphism varies among populations. For any trait with such an interaction (standard length, head length, caudal peduncle depth, and length of gill rakers), we quantified dimorphism within each population separately by calculating between-sex t-statistics.

### Trait-environment correlations

We tested whether our estimates of population trait means, dimorphisms, and allometries, were related to among-lake differences in abiotic and biotic environmental variables. Such correlations among lakes are typically interpreted as evidence of adaptive value. We tested whether each stickleback trait (lake means, dimorphism, or allometric slope) was correlated with many environmental traits. All lake traits involving size or distance were log transformed. Area is frequently mentioned as a key variable distinguishing relatively benthic versus limnetic lake populations (e.g. Bolnick and Ballare 2020). We also tested for relationships with *S. solidus* prevalence (which we expect to increase with reliance on limnetic prey) and lake elevation.

For a more comprehensive approach to evaluating the adaptive basis of heritable morphology, we used a machine learning approach to contrast stickleback morphology with 276 landscape, climate, water column, and zooplankton community predictors. Environmental data processing steps are described in depth in Supplement 1 (Methods for Environmental Covariates). Supplementary Figure S2 contains ecosystem gradient profiles across study lakes. Given the high-dimensional, correlated nature of our data, we conducted a non-parametric variable selection workflow by fitting independent Random Forest models for each morphometric trait using the ranger function (1,000 trees, *mtry* = *N_vars_*/3; Wright and Ziegler, 2017; Breiman, 2001). Top features surpassing a dynamic permutation importance threshold of *100/N_vars_* were selected for confirmatory linear regressions using the lm function (R Core Team, 2026). Prior to model fitting, continuous predictors were *z*-score standardized to derive standardized effect sizes (*β*). Highly influential observations exceeding a Cook’s Distance threshold of *4/N* were systematically removed to prevent outlier bias (Bollen and Jackman, 1985). Finally, to complement these univariate models and evaluate the global, multi-trait influence of zooplankton gradients, we performed a distance-based Redundancy Analysis (dbRDA) via the *dbrda* function on a multivariate Euclidean phenotypic distance matrix, calculating explained variance via the *RsquareAdj* function and testing overall model significance and individual axis margins using 999 random permutations via the *anova.cca* function (Oksanen et al., 2022).

## Results

### Among-population morphological variation in a common garden setting

Multivariate analysis of size-adjusted traits revealed significant differences among populations reared in the lab environment (Figure 1; MANOVA Pillai’s trace = 4.0744, *F*_364_, _5852_ = 6.60, *p* < 0.001). There is appreciable overlap among population trait distributions along the two leading discriminant axes (Figure 1A), but the means are clearly distinct (Figure 1B). These leading two axes explain 34% and 18% of overall phenotypic variance, and are dominated by body depth and standard length respectively. The top 11 discriminant axes all show significant among-population variation (Supplementary Table S3), suggesting that heritable phenotypic divergence is highly multivariate rather than being confined to a single major benthic-limnetic axis. Using a confusion matrix, the LDA correctly classified 70.79% of individuals to their population of origin (Supplementary Table S4). With 27 populations, we would expect 1/27 (3.7%) success rate if each individual was randomly assigned.

**Figure 1:**
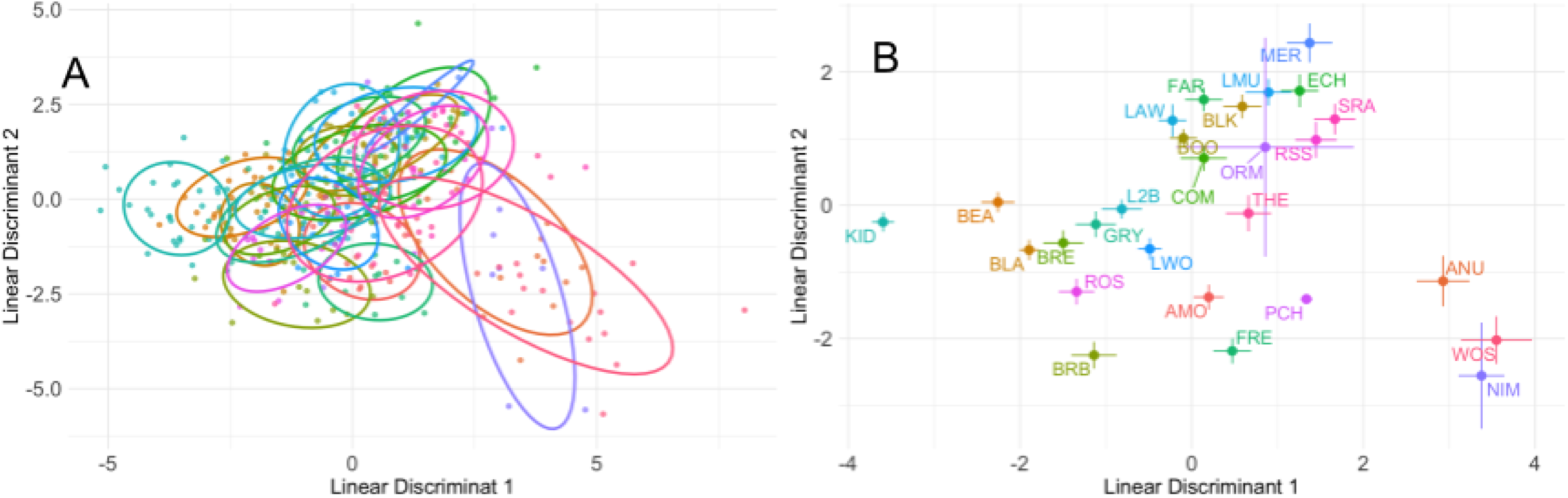
Linear discriminant analysis (LDA) of size-adjusted morphological traits across populations. A) Each point represents an individual, colored by population. Ellipses indicate the 68% density regions (approximately one standard deviation) for each population, illustrating the distribution and overlap of lab-raised populations in multivariate trait space. Separation among ellipses reflects the degree of morphological differentiation among populations based on size-corrected traits. B) Population-level means of linear discriminant scores (LD1 and LD2) derived from size-adjusted morphological traits. Points represent population means, and error bars indicate +/− 1 standard error for each axis. Supplementary Table S3 has the trait loadings on the LD axes. Supplementary Table S4 contains the confusion matrix obtained from the LDA. Lake name abbreviations are provided in (B), with full names provided in Supplementary Table S1.

Univariate ANOVAs confirm that all 14 measured traits exhibited significant among-population differences after Sequential Bonferroni correction (Supplementary Table S5). Here we highlight results for a few select traits to illustrate the broader trend. The mean population number of armor plates ranged from 7.11 to 14.6, with the lowest counts in Kidney and Beaver Lakes and highest in Woss, Anutz, and Nimpkish Lakes (Figure 2A). The mean number of gill rakers ranged between 17.8 and 21.7, lowest in Kidney and Beaver Lakes, highest in Merrill Lake (Figure 2B). Size-adjusted gill raker length (Figure 2C) and body depth (Figure 2D) were also leading sources of variation.

**Figure 2:**
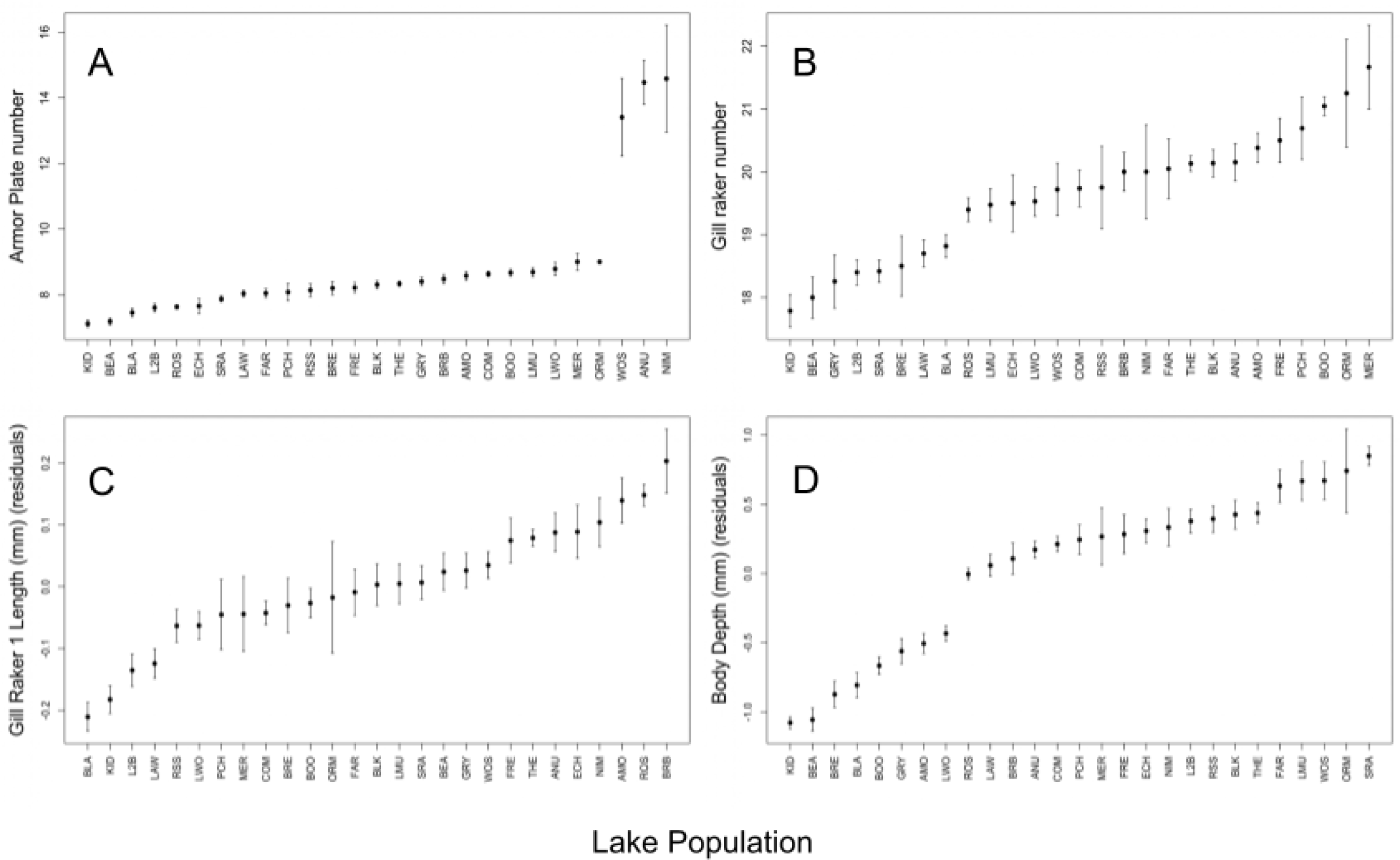
Mean and +/− 1 standard error confidence intervals of armor plate number. (Panel A), gill raker number (Panel B), size-adjusted length of the longest gill raker (mm) (Panel C), and size-adjusted body depth (mm) (Panel D) of lab-raised populations from 27 lakes on Vancouver Island. Lakes are ordered along the x axis by trait value. The full lake names are given in Supplementary Table S1.

Although all univariate linear measurements were correlated with body length, for a few traits this allometric relationship varied among populations (Supplementary Table S5). Head length (*F* = 2.26, *p* = 0.0004) and snout length (*F* = 2.19, *p* < 0.0006) have a significant population x length interaction after Sequential Bonferroni correction. Three other traits were significant prior to multiple test correction: eye diameter (*F* = 1.61, *p* = 0.0285), caudal peduncle depth (*F* = 1.69, *p* = 0.0180), and body depth (*F* = 1.69, *p* = 0.0177). Thus, prior to Bonferroni correction, 5/14 traits (36%) showed allometric variation at α = 0.05 cutoff.

### Among-trait correlations across populations

We find numerous pairs of traits whose means covary among populations reared in the lab (Figure 3; statistical results in Supplementary Table S6). Examples include a positive correlation between snout length and head length (r = 0.86, *p* < 0.001) and between caudal peduncle and gill raker density (r = 0.62, *p* < 0.001). 19 out of 91 pairwise correlations (21%) are significant at α = 0.05. Some of the uncorrelated comparisons are noteworthy for being surprising, such as a lack of relationship between gill raker length versus gill raker number, both canonical traits in the benthic-limnetic axis of divergence (Schluter and McPhail, 1992; MacColl, 2009).

**Figure 3:**
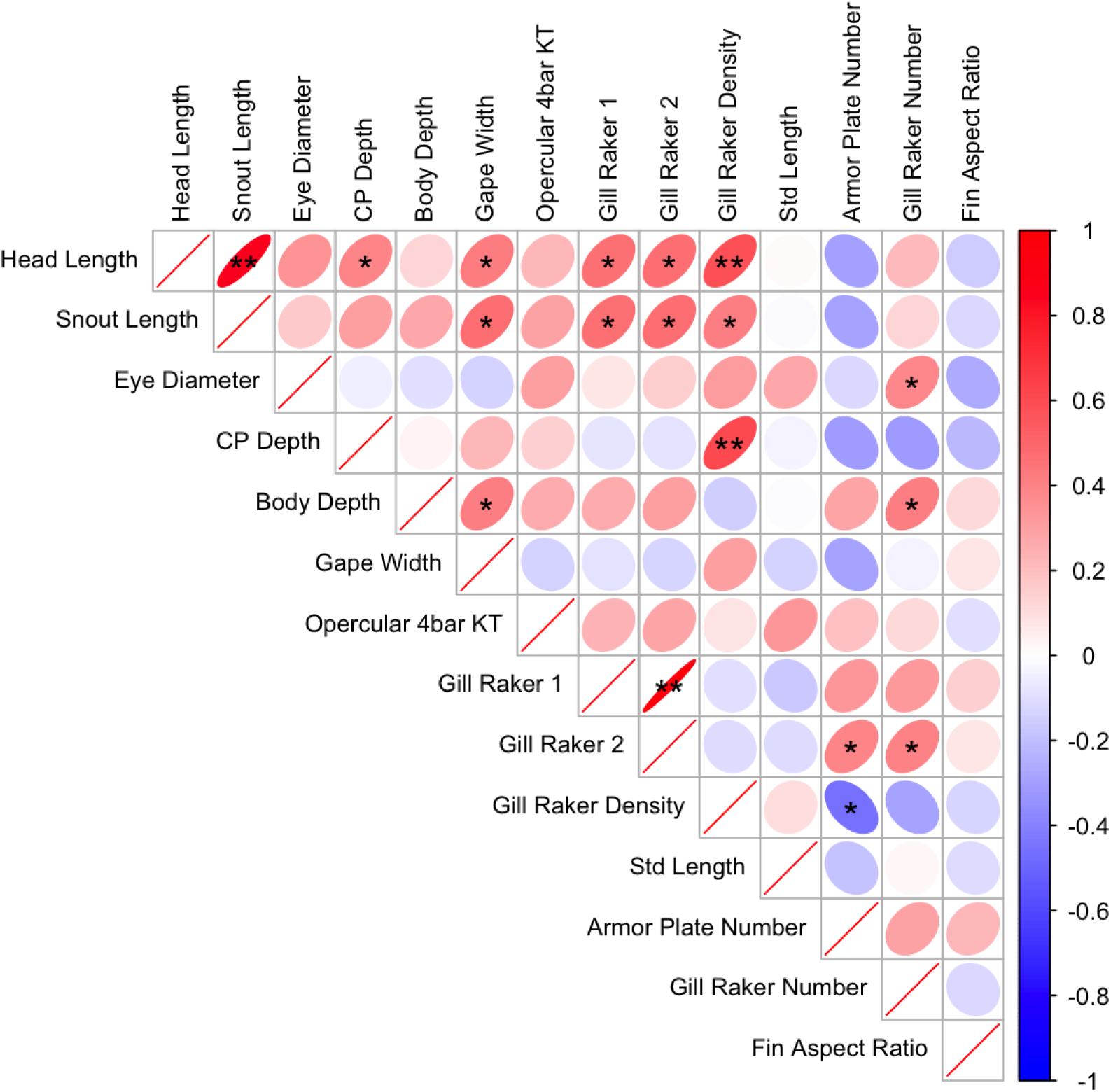
Correlation between the 14 morphological traits’ population means. Ellipses indicate the strength and direction of Pearson correlations, with red representing positive relationships and blue representing negative relationships; color intensity and ellipse shape reflect correlation magnitude. An asterisk (*) in an ellipse signifies significance (*P* < 0.05) while double asterisks (**) signifies significance after Sequential Bonferroni correction. The statistical test details (r and *p* values) are in Supplementary Table S6.

**Figure 4:**
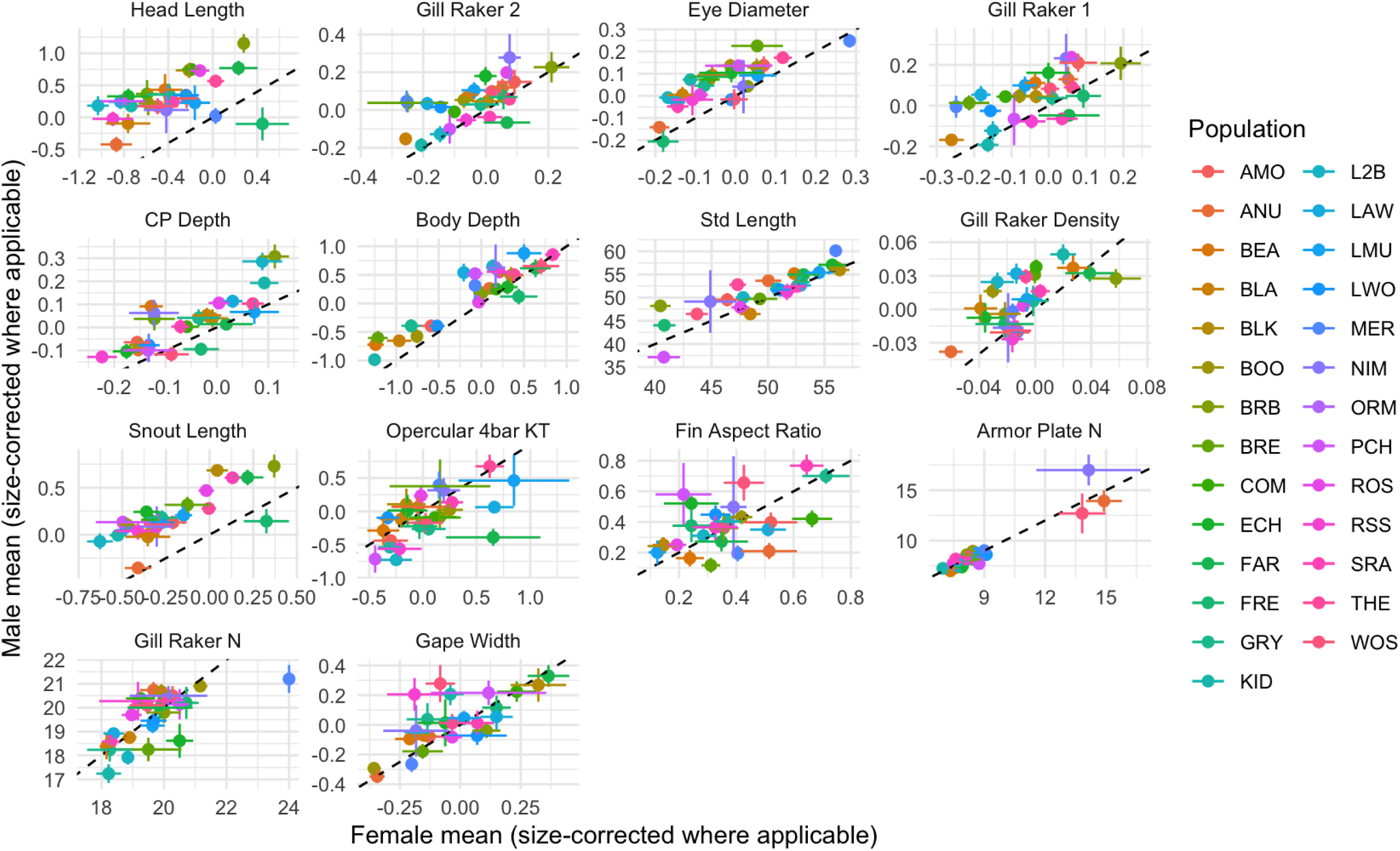
Sexual dimorphism across morphological traits, ordered from most to least dimorphic (top left to bottom right) based on the magnitude of difference between male and female mean trait values averaged among lakes. Opercular 4-bar KT, fin aspect ratio, armor plate number, gill raker number, and gape width showed no SD. All traits are size-adjusted for standard length where applicable, except for standard length, armor plate number, and gill raker number. Each point represents a population, abbreviated lake names are explained in Supplementary Table S1. The dashed 1:1 line indicates no sexual dimorphism; deviations from this line reflect the extent and direction of sex differences. Error bars represent +/− 1 standard error. Statistical results of ANOVAs with lake, sex, and lake x sex interaction effects are given in Supplementary Table S5.

### Sexual dimorphism is seen in most traits

Dimorphism was observed in 9 of the 14 morphological traits after size standardization, p < 0.001 (Supplementary Table S5). The non-dimorphic exceptions included gape width (F = 1.75, p = 0.19), armor plate number (F = 0.02, p = 0.9131), gill raker number (F = 0.31, p = 0.5754), opercular 4-bar KT (F = 2.85, p = 0.0919), and fin aspect ratio (F = 1.17, p = 0.2801). Across sexually dimorphic traits, males generally exhibited larger size-adjusted trait values than females, although the magnitude and direction of dimorphism varied among traits and populations (Figure 4).

The magnitude of sexual dimorphism was mostly similar across lab-raised populations. That is, we observed non-significant (p > 0.05) population x sex interactions for 9 of the 14 traits (Supplementary Table S5). The exceptions were head length (F = 1.65, p = 0.024), caudal peduncle depth (F = 1.78, p = 0.0113), gill raker 1 length (F = 1.96, p = 0.0042), gill raker 2 length (F = 1.96, p = 0.0416) and standard length (F = 2.25, p = 0.0005, the only population x sex interaction to survive Bonferroni correction).

### Correlations with environmental covariates

For morphological traits, lab-reared population means were correlated with certain environmental features of their parents’ lakes (Figure 5; Supplementary Table S7). Armor plate number was correlated with lake area (Figure 6A), perimeter (r = 0.64, *p* < 0.001), and maximum depth (r = 0.73, *p* < 0.0001). Elevation is negatively correlated with armor plate number (r = –0.44, *p* = 0.0248). Caudal peduncle depth is negatively correlated with lake area (Figure 6B), perimeter (r = –0.48, *p* = 0.0122), and distance away from the ocean (r = –0.47, *p* = 0.0146). Notably, two standard features of stickleback ecotypes, gill raker number (Figure 6C) and gill raker length (Figure 6D), show no meaningful correlation with the expected lake traits. Lake elevation was positively correlated with fish standard length (Figure 6E) and eye diameter (r = 0.52, *p* = 0.0066). Distance from the ocean is also correlated with standard length and eye diameter.

**Figure 5:**
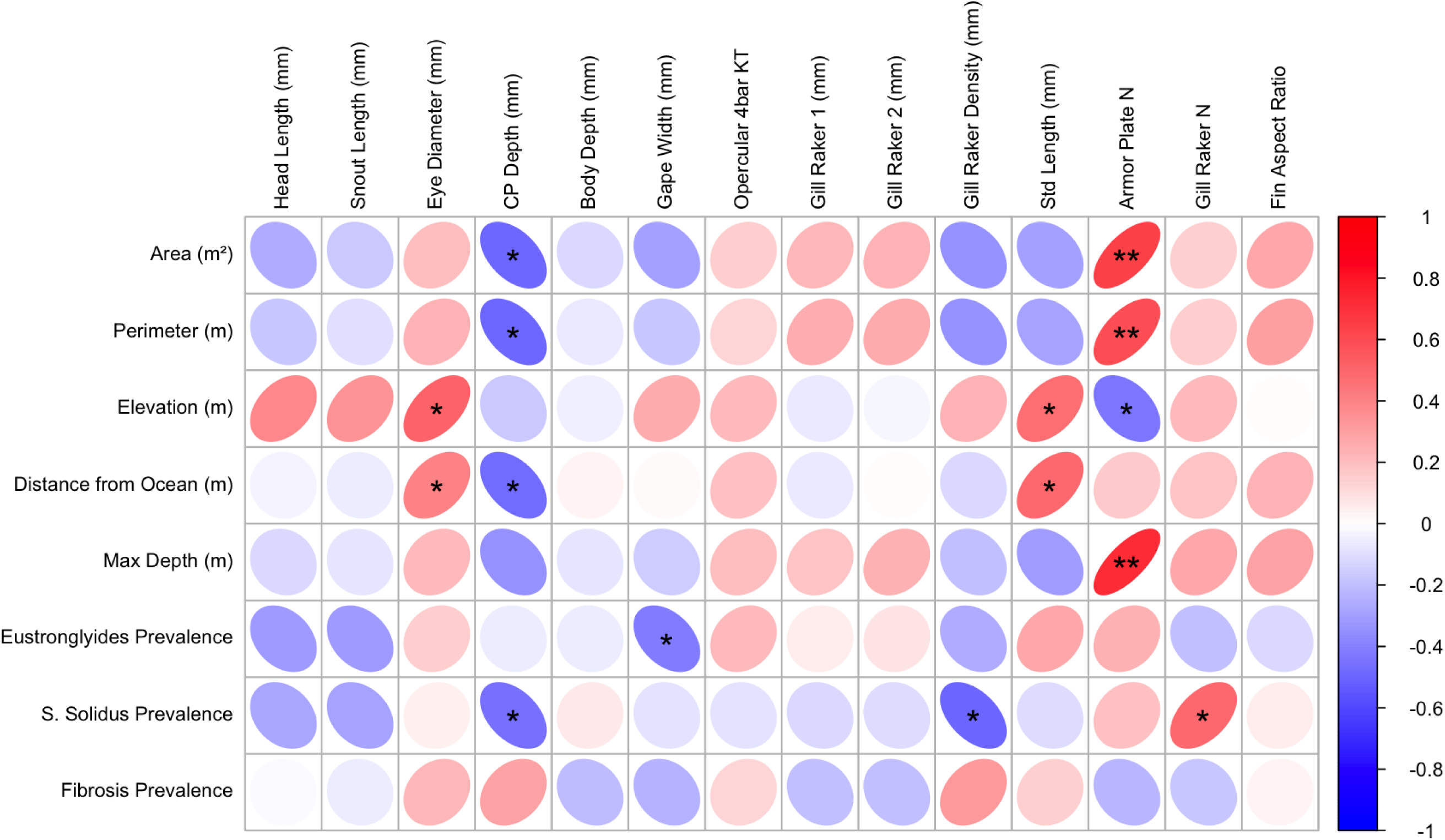
Correlations between environmental variables (lake characteristics and population parasite prevalence) versus size adjusted population mean morphological traits measured on lab-raised fish. Ellipses indicate the strength and direction of Pearson correlations, with red representing positive relationships and blue representing negative relationships; color intensity and ellipse shape reflect correlation magnitude. An asterisk (*) in an ellipse signifies significance (*p* < 0.05) while double asterisks (**) signifies significance in terms of sequential Bonferroni. Statistical effect sizes and *p* values are presented in Supplementary Table S8.

**Figure 6:**
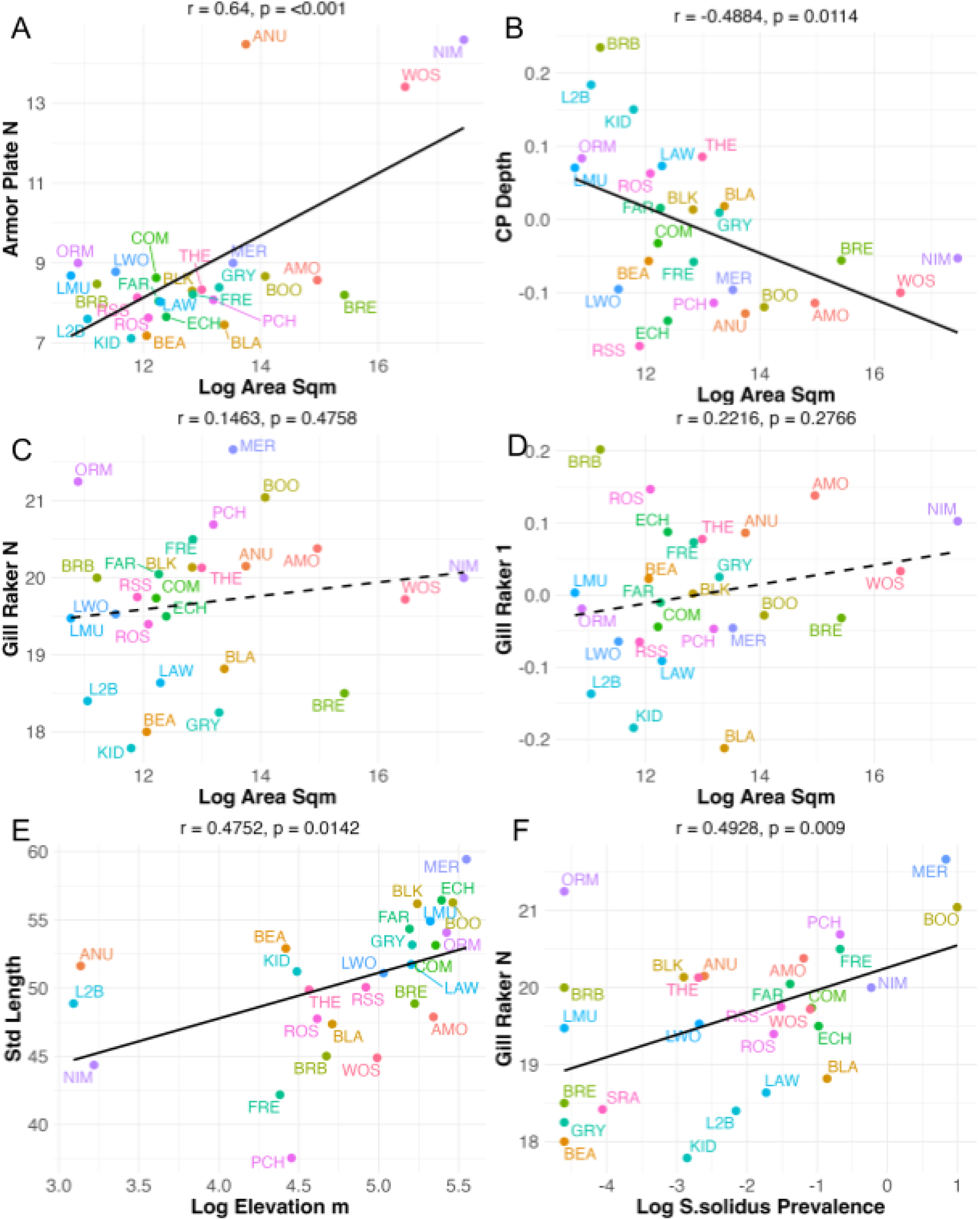
Trait-environment correlations between. (A) log area of lake (m^2^) and population mean armor plate number; (B) area of lake (m^2^) and mean size-adjusted caudal peduncle depth (mm); (C) area of lake (m^2^) and population mean gill raker number; (D) area of lake (m^2^) and maximum gill raker length; (E) elevation (m) and mean standard length (mm); and (F) *S. solidus* prevalence and mean gill raker number. Correlation coefficients and corresponding *p* values are listed above each panel. Supplementary Table S8 contains all correlation coefficients and *p* values.

The prevalence of wild *S. solidus* tapeworms was positively correlated with gill raker number (Figure 6F) and negatively correlated with caudal peduncle depth (r = –0.46, *p* = 0.0185) and gill raker density (r = –0.50, *p* = 0.0086) in the lab. These trends are consistent with expectations for limnetic ecomorphology conferring greater *S. solidus* exposure (Fleischer et al., 2021). We found no significant morphological covariates with fibrosis severity in the field, an immune trait involved in resistance to *S. solidus*. Environmental features of lakes did not explain any of the variation in trait allometry (population-specific trait-length slopes). Nor was the magnitude of heritable dimorphism (population-specific t-values) correlated with lake traits.

In the multivariate machine learning analysis, our evaluations reveal that stickleback morphological traits are primarily shaped by macro-scale landscape and climate characteristics rather than localized water column properties or dietary shifts. Across individual feature models (Figure 7, Supplementary Table S8), physical geography and regional climate metrics emerged as the most robust drivers of phenotypic variance. Specifically, lake perimeter strongly predicted armor plate counts (b = 1.532, SE = 0.156, t = 9.83, *p* < 0.001), as did maximum estimated lake depth (b = 0.993, SE = 0.141, t = 7.02, p < 0.001). In contrast, localized water column parameters consistently demonstrated weaker or non-significant associations with individual traits. For example, while variation in dissolved oxygen (SD_DO_Percent) showed a minor association with caudal peduncle depth (b = 0.047, SE = 0.016, t = 3.00, *p* = 0.007), key morphological traits such as fin aspect ratio and gape width lacked any statistically significant predictors across potential environmental drivers. Appendix Table 1 provides information on covariate names, data sources, and ecological context. Supplementary Figure S2 provides values of each environmental metric for all lakes.

**Figure 7:**
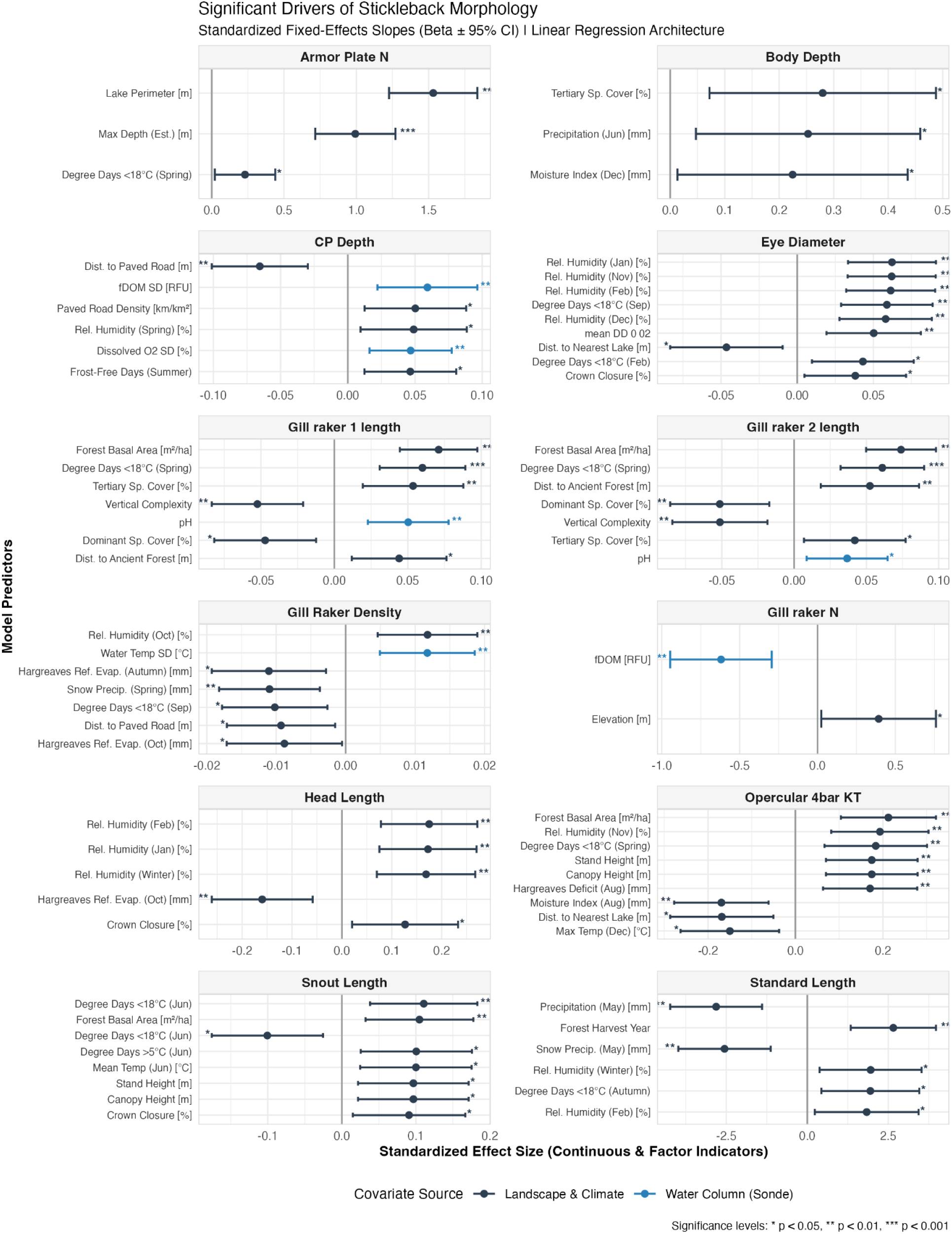
Significant environmental drivers of stickleback morphology. Standardized regression coefficients (*β* ± 95% confidence intervals) from linear models relating lake environmental covariates to population mean morphological traits of common garden-reared threespine stickleback. Each panel represents a separate morphological trait, and only environmental variables with statistically significant relationships are shown. Positive coefficients indicate that the trait increases with increasing values of the environmental covariate, whereas negative coefficients indicate the opposite relationship. Black points represent landscape and climate variables, and blue points represent water chemistry variables measured with a YSI sonde. Horizontal bars indicate 95% confidence intervals.

Furthermore, multivariate zooplankton community analysis confirms that localized food-web structure has a negligible effect on stickleback morphology across lakes. Although the Hellinger-standardized Principal Components Analysis (PCA) successfully captured localized zooplankton community variation, with the first two axes accounting for 33.3% (PC1) and 14.0% (PC2) of the food-web variance, these community gradients did not map onto morphological divergence. The distance-based Redundancy Analysis (dbRDA) confirmed that neither the overall multivariate model (F = 1.0362, *p* = 0.398, R^2^ = 0.0827, Adjusted R^2^ = 0.0029; Supplementary Table S9A and Supplementary Table S9B) nor the individual marginal axes (PC1: F = 0.9971, p = 0.440; PC2: F = 1.0268, p = 0.415; Table S9C) explained significant morphological variation. Collectively, these results indicate that stickleback morphology is predominantly driven by broad, physical landscape characteristics and regional climate regimes rather than spatial variation in water chemistry or zooplankton community composition.

## Discussion

Distinguishing phenotypic variation that reflects heritable genetic divergence from variation by phenotypic plasticity is a central challenge for evolutionary biologists. Many studies identify strong trait-environment correlations in natural populations, but these patterns alone cannot distinguish whether populations have evolved genetically or whether individuals simply develop different phenotypes when exposed to different environments. Common garden experiments provide a powerful approach for separating these alternatives as they minimize environmental differences while preserving genetic differences among populations. Common gardens cannot completely exclude non-genetic inherited effects, such as maternal effects (English et al, 2015) or epigenetic inheritance (Whipple and Holeski, 2016), but they reduce environmental sources of phenotypic variation, which allows heritable differences among populations to be identified with much greater confidence. Our study extends this approach by examining an unusually large number of populations, allowing us to evaluate not only whether morphological divergence is heritable, but also whether the ecological patterns observed in wild populations persist when environmental effects are minimized.

Many previous studies of stickleback have documented that morphological traits differ repeatedly among populations spanning different habitats (McPhail, 1992; Cresko et al., 2004). Examples include marine versus freshwater (Reimchen, 1983; Bell and Foster, 1994; McKinnon and Rundle, 2002; Colosimo et al., 2005), lake versus stream (Hendry and Taylor, 2004; Kaeuffer et al., 2012; Oke et al., 2016; Stuart et al., 2017), and benthic versus limnetic lake habitats (Schluter and McPhail, 1992; MacColl, 2009; McGee et al,. 2013; Dean et al., 2023). However, these previous studies generally used wild-caught stickleback, limiting our ability to determine whether these trait-environment associations represent evolved by genetic divergence or by environmentally induced plasticity. Additional studies have used common garden rearing experiments to confirm that between-population differences are partly heritable (Supplementary Table S1), but these common garden studies typically consider only a few populations, so they cannot effectively test whether trait-environment correlations reflect adaptive genetic variation.

Here, we use a common garden experiment with 27 stickleback populations to show that among-population trait variation is heritable. Though we were unable to replicate some classical trait-environment correlations from prior stickleback studies (i.e., gill raker traits, body depth), some population traits we measured were partly correlated with source lake environments, suggesting a role of adaptive evolution.

### Every morphological trait differs among populations

All 14 measured morphological traits varied among the 27 populations reared under common garden conditions, providing strong evidence that morphological variation among these stickleback populations has a substantial heritable component. Among-lake variation in stickleback is often framed in terms of a univariate axis from relatively benthic to limnetic ecotypes (Bentzen and McPhail, 1984; Schluter and McPhail, 1992). However, the leading discriminant axis only accounted for 34% of morphological variation among individuals in our common garden. Rather than just one major axis, we observed statistically significant differences among populations for the top 11 discriminant axes. Thus, heritable among-lake differences are highly multivariate. The accuracy was relatively strong at 70.79%, substantially higher than if they were randomly assigned (3.7%). This demonstrates that populations possess distinct multivariate morphological signatures despite considerable overlap among individuals. Because these fish were reared under common garden conditions, these extensive morphological differences cannot be attributed solely to differences in developmental environment. Instead, these results demonstrate that substantial genetic variation exists among populations, providing the raw material upon which natural selection can act.

Most univariate traits exhibit allometric increases with body length (except gill raker number and armor plate number. The slopes of these allometric relationships were generally similar across populations, with the exception of five traits; head length, snout length, eye diameter, caudal peduncle depth, and body depth all exhibit some variation in allometric slopes among populations. This variation in trait development may reflect differences in populations’ ecology. For example, eye diameter is associated with visual performance in detecting predators and finding prey (Beston & Walsh, 2019; Hall & Ross, 2007; Land & Nilsson, 2012). Larger eyes also allow clear visualization in darker waters (Andersson et al., 2024). Thus, differences in scaling may indicate adaptations to varying light environments or habitat types. Similarly, variation in the scaling of head length and snout length suggests differences in feeding mechanics among populations, as these traits are associated with suction feeding performance, particularly in limnetic environments where sticklebacks capture evasive copepod prey (McGee et al., 2013).

We observed a moderate number of correlations between trait means. For example, caudal peduncle depth and gill raker density are strongly positively correlated across populations. Such correlations among traits are expected for either of two reasons. First, sets of traits may be genetically linked, so that selection acting on one trait’s mean could consistently produce correlated changes in a second trait, resulting in among-lake correlations between trait means. This is almost certainly the case for the positive correlation between snout length and head length, which measure closely interrelated skeletal features. Alternatively, two or more traits may experience correlated selection: environments that favor an increase in one trait mean might consistently favor an increase in the mean of a second trait as well. This may be the case for relationships between structurally distinct traits such as caudal peduncle depth and gill raker density. However, our study is not designed to distinguish between effects of G matrix structure, versus correlated selection.

### Sexual dimorphism occurs yet differs among populations

Sexual dimorphism in morphological traits has been widely documented in stickleback populations, using wild-caught fish (Reimchen and Nosil, 2004; Aguirre et al., 2008; Spoljaric and Reimchen, 2008; Aguirre and Akinpelu, 2010; Leinonen et al., 2011; McGee and Wainwright, 2013), but some of these between-sex differences may be a result of divergent phenotypic plasticity acting on sexes. Or, sexually antagonistic selection could exaggerate sex differences if natural selection tends to remove individuals which phenotypically most resemble the opposite sex. By rearing fish in a common garden setting we are able to minimize both selective forces, thereby confirming that there is a genetic basis to sexual dimorphism in this system. Dimorphism was observed in most morphological traits, with the exception of gape width, armor plate number, gill raker number, and opercular 4-bar KT. For traits exhibiting dimorphism, males generally had larger size-adjusted trait values than females. Consistent with our results, Leionen et al., (2006) found that male stickleback also had larger body sizes than females with no differences in armor plate number. Importantly, the magnitude of sexual dimorphism did not vary noticeably among lake populations (no population x sex interaction) for most of the traits we measured. This suggests that the genetic and developmental mechanisms underlying sexual dimorphism are largely conserved among populations for these traits. Some of these instances of conserved dimorphism are surprising; for example, previous studies that include lakes we examined in this study have found the number of gill rakers to be sexually dimorphic, and differences in dimorphism among lakes (Bolnick and Lau 2008). Our results suggest that the previously documented gill raker dimorphism may be a result of within-generation processes (development or selection) acting in wild fish. This is why it may be critical to use common garden experiments to study sexual dimorphism. We do observe a few traits with population x sex interactions (standard length, head length, caudal peduncle depth, and gill raker lengths), suggesting that some aspects of morphological dimorphism are actively diverging among populations.

### Trait-environment correlations (and the lack thereof)

Correlations between environmental variables and phenotypic traits are commonly interpreted as evidence for adaptive divergence in response to natural selection. Many studies have documented such trait-environment correlations across stickleback populations. But, most are based on wild-caught fish, unable to rule out the added contribution of plasticity. Or, they use too few lab-reared populations to have much statistical power. Our study is able to test for evolved correlations using an unusually large number of lab-reared populations. We document correlations between multiple sets of limnological variables and stickleback traits that are likely to reflect adaptive evolution.

Firstly, we found that lake surface area is positively related to armor plate number. Stickleback that are in environments with more predatory fish have more armor plates to protect from predation (Hagen & Gilbertson, 1972; Moddie & Reimchen, 1976; Bell & Richkind, 1981; Kitano et al., 2008); the three largest lakes that drive this correlation may have an especially intense risk of predation by large salmonid fish. A caveat is that these three lakes are all in the same watershed, so may not be fully independent from a perspective of ancestry. The lowest lake in the watershed (Nimpkish) likely has recurrent gene flow from marine fish sustaining high armor. Lake size (both area and perimeter) is also correlated (negatively) with caudal peduncle width. Smaller caudal peduncles are associated with energy-efficient swimming (Martinez et al., 2021), which may be helpful in more pelagic foraging in large lakes. Machine learning analyses revealed that morphological traits were heavily associated with climatic variables such as water temperature, moisture deficit, humidity, and forest characteristics surrounding the lakes, instead of the generally hypothesized lake size. For example, warmer temperatures and greater forest cover were generally associated with larger values of several morphological traits, including head length and gill raker length, whereas greater humidity was positively associated with eye diameter and several other traits. Gill raker traits showed additional associations with landscape characteristics. In contrast, gill raker traits decreased with increasing dominant tree cover and human landscape. These results suggest that broad climatic and surrounding landscape conditions may contribute to morphological divergence among populations in ways that extend beyond the traditional focus on lake size and benthic-limnetic ecology. These results suggest that morphological divergence among populations is influenced by large-scale abiotic environmental gradients, which have not previously been identified as a major driver of adaptive divergence among lake stickleback. The precise biological mechanisms generating this divergent natural selection among lakes remain to be determined. But, these trait-environment correlations do not support the long-standing view that lake size (or, similarly, relative littoral area) is the primary driver of a benthic-limnetic axis of ecotype variation, acting via the relative availability of planktonic zooplankton versus benthic insect larvae.

Lake elevation was positively correlated with standard length and eye diameter, while being negatively correlated with armor plate. These results might reflect selection due to climatic variables associated with elevation, but the lakes we sampled span a small elevation range (22 to 257 meters). Alternatively, elevation range can be related to filters in the original upstream colonization process, if these traits affected sticklebacks’ capacity for up-river dispersal. Consistent with this colonization filter, lake distance from the ocean is positively correlated with standard length and eye diameter and negatively correlated with caudal peduncle depth. Smaller caudal peduncles would likely have allowed individuals to efficiently swim farther inland.

We also found some surprising negative results. Gill raker number and length are typically correlated with lake size, being classic features separating benthic and limnetic populations (Schluter and McPhail, 1992; Kristjánsson, 2005; MacColl, 2009; Wund et al., 2012). Body depth and gape width are also widely reported to distinguish benthic versus limnetic ecotypes. However, although gill raker traits varied among lab-raised populations, no gill raker measures were correlated with lake area, depth, or other limnological features. Body depth and gape width are also uncorrelated with lake area or depth. We found no evidence that differences in zooplankton community composition and density explained among-population morphological divergence. Although PCA summarized substantial variation in zooplankton communities among lakes, neither the overall dbRDA nor the individual community axes significantly explained morphological variation. These negative results are contrary to our expectations from previous studies (including some from these same lakes). Admittedly, we cannot rule out the possibility that weak correlations exist: with 27 lakes our statistical power is good, but not perfect. Regardless, our results do suggest that the classic benthic-limnetic axis or zooplankton community that is widely documented in stickleback may not persist in laboratory settings. The implication is that this ecotypic variation among allopatric lakes may have a strong contribution from phenotypic plasticity.

We do, however, find a relationship between gill raker traits and a biotic variable. The existing stickleback literature leads us to expect that more limnetic stickleback populations have more gill rakers, as an adaptation to consume more zooplankton including copepods (Schluter & McPhail, 1992; Bolnick and Ballare 2020). Consequently, limnetic populations should have a higher rate of *S. solidus* exposure and higher prevalence, all else being equal (Stutz and Bolnick 2014; Weber et al. 2022). We find a significant positive relationship between *S. solidus* prevalence (in wild-caught samples) and gill raker number in lab-raised fish. There is a corresponding negative relationship with gill raker density. *S. solidus* prevalence was also negatively correlated with caudal peduncle depth: the smaller peduncle is also typical of a more fusiform limnetic body shape for mid-water foraging. These correlations indicate that evolved morphological trait differences among lake populations affect sticklebacks’ risk of infection by trophically transmitted parasites. The main immune defense that stickleback evolve against *S. solidus* infection is a peritoneal fibrosis response that can greatly suppress tapeworm growth and viability (Weber et al. 2022). We therefore expected to find a positive relationship between fibrosis severity and ecomorphology, which was not found. This is not entirely surprising given that observed rates of fibrosis in the wild are dependent on genetic variation in baseline fibrosis, fibrosis response to tapeworm exposure, and exposure rates.

More generally, these findings illustrate why common garden experiments remain essential for understanding adaptive evolution. Trait-environment associations measured in wild populations are frequently interpreted as evidence of local adaptation, yet they can arise from plastic developmental responses to local environments. By separating genetic from environmental sources of phenotypic variation, common garden experiments allow stronger inference about which traits have evolved through natural selection and which primarily reflect developmental plasticity. Our results suggest that some widely accepted stickleback ecotype differences may owe a greater contribution to plasticity than previously appreciated, whereas other traits retain clear signatures of heritable adaptive divergence.

## Conclusion

Our study demonstrates that there is substantial multivariate morphological variation among 27 lake populations of threespine stickleback. This supports the prevailing view that much of the observed phenotypic divergence among wild stickleback populations has a genetic basis. Some of the heritable trait differences are associated with lake geography (surface area, depth, elevation), or with climatic variables. These trait-environment correlations are consistent with adaptive evolutionary divergence driven by natural selection. However, this variation does not fall along the major benthic-limnetic axis. And, some of the classic trait-environment correlations in stickleback are not found in our data. The often-described benthic-limnetic variation led us to expect correlations between lake size versus gill raker length, number, gape width, and body depth, but these were not confirmed by our data from common garden fish. We also confirmed that sexual dimorphism has a substantial heritable basis, with consistent differences between males and females across most populations, indicating that the developmental mechanisms underlying dimorphism are largely conserved despite extensive population divergence. These results reiterate the importance of using common garden rearing experiments with a large sample of populations, to re-evaluate potentially adaptive trait-environment correlations seen with wild-sampled individuals. Additionally, they suggest that lake microclimate variables may be a majorly overlooked driver of adaptive morphological variation in stickleback.

Furthermore, our findings highlight the importance of distinguishing heritable evolutionary divergence from environmentally induced phenotypic plasticity when interpreting phenotypic variation in natural populations. Although trait-environment correlations are often used as evidence for local adaptation, our results show that some widely accepted associations weaken or disappear when environmental effects during development are controlled. Common garden experiments therefore remain an essential approach for identifying which phenotypic differences represent evolved genetic divergence and which primarily reflect plastic responses to local environments. Understanding the relative contributions of heredity and plasticity will be increasingly important for predicting how populations adapt to environmental change.

## Supplementary Figures

**Supplementary Figure S1:**
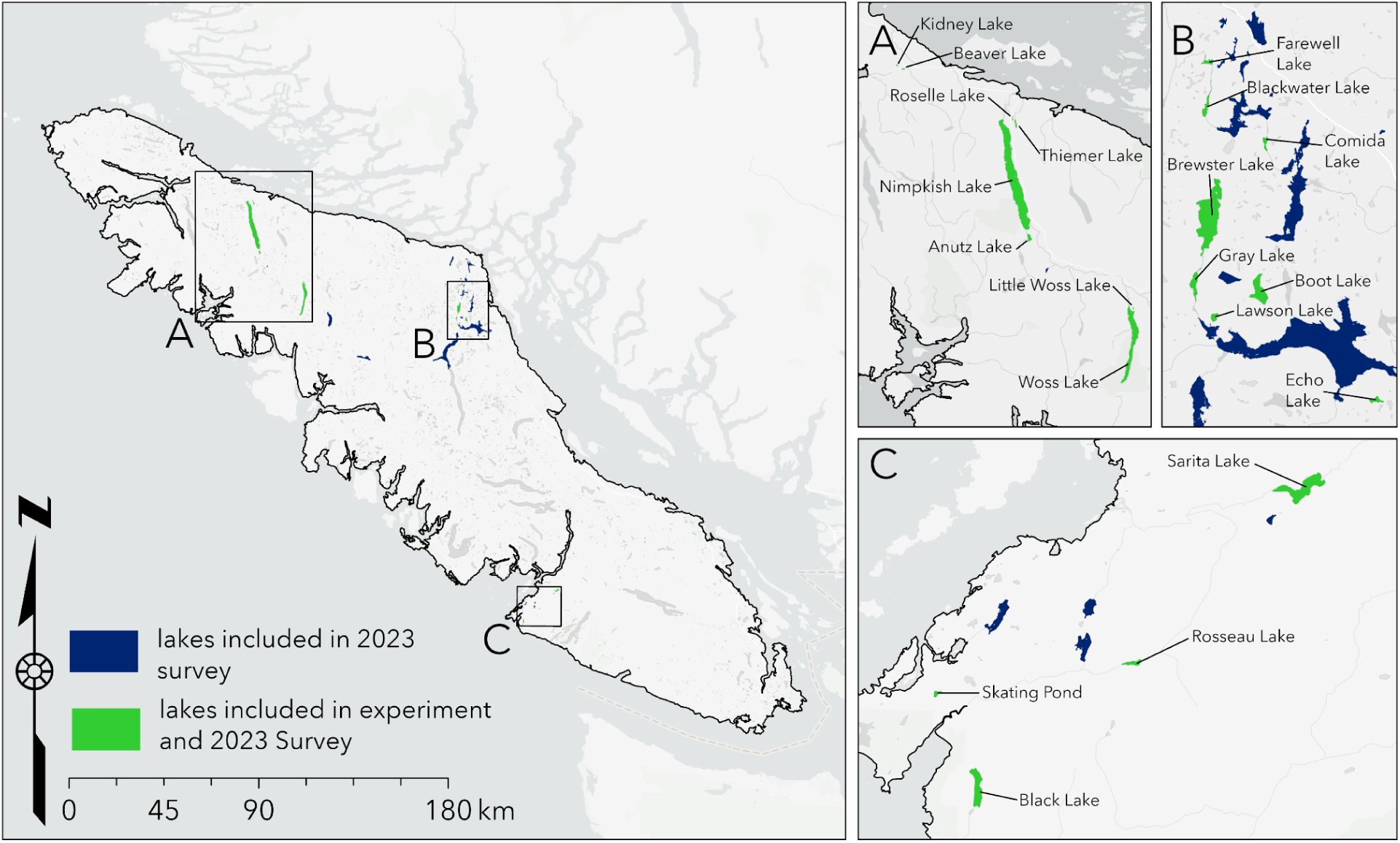
Map of location of lakes.

**Supplementary Figure S2:**
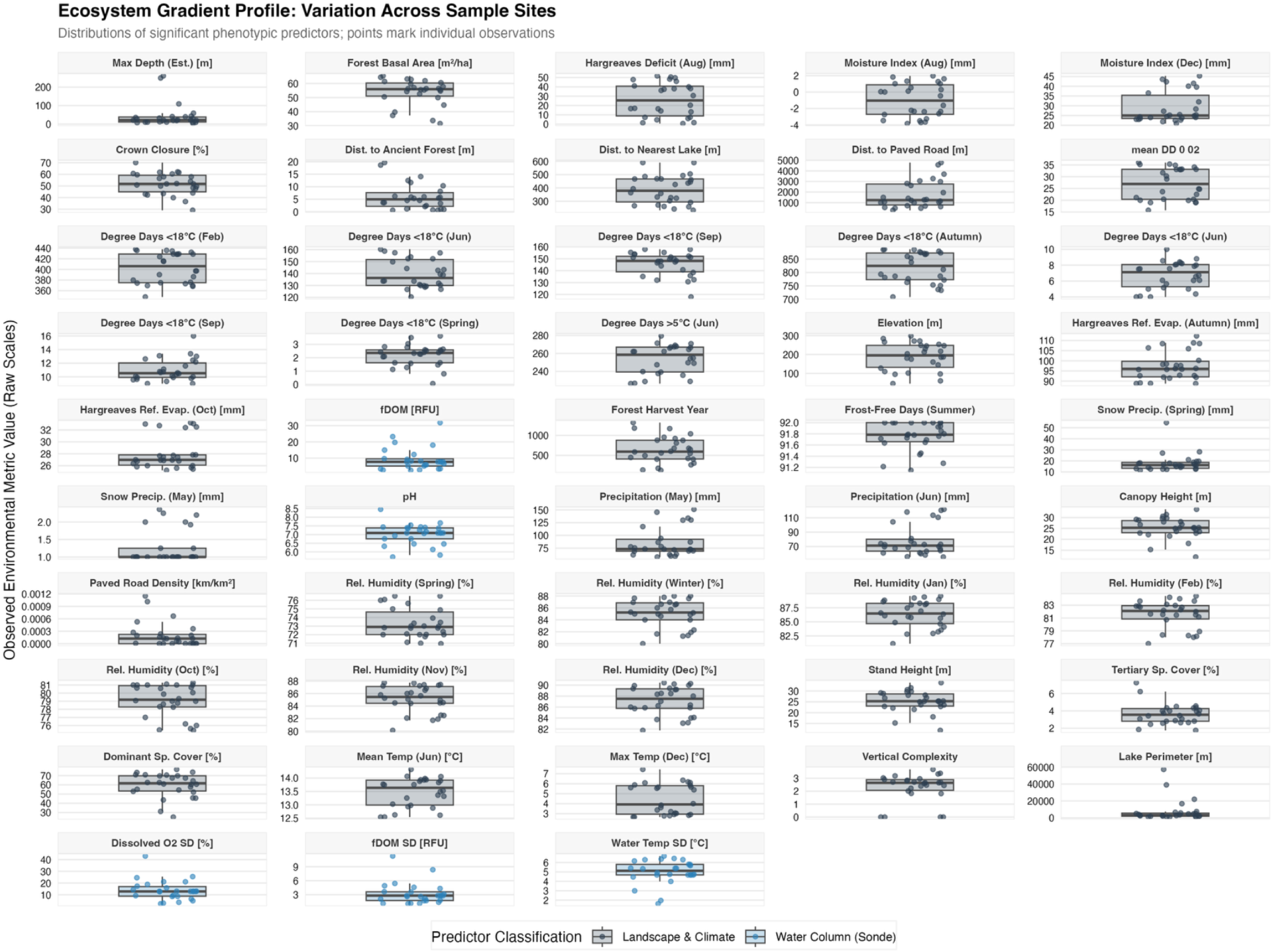
Ecosystem gradient profiles across study lakes. Distribution of environmental predictor variables measured across the 27 study lakes. Each panel shows a boxplot summarizing the distribution of a single environmental variable, with points representing individual lake values. Variables include landscape and climate characteristics (gray) and water-column (Sonde) measurements (blue). These predictors were evaluated as potential environmental correlates of morphological variation among common-garden reared threespine stickleback populations. The figure illustrates the range and variability of environmental conditions represented by the sampled lakes and provides context for subsequent analyses relating environmental gradients to morphological divergence.

## Supplementary Tables

**Supplementary Table S1:** Stickleback studies involving common-garden experiments studying morphology.

| **Paper** | **Ecotype** | **# of Populations Reared** |
| --- | --- | --- |
| Lavin and McPhail, 1993 | Lake-stream | 3 populations (1 lake and 2 streams) |
| Day et al., 1994 | Benthic-limnetic diet | 4 populations (benthic worms and plankton, limnetic worms and plankton) |
| Hatfield, 1997 | Benthic-limnetic | 6 populations (limnetic, limnetic backcross, F1 hybrid, F2 hybrid, benthic, benthic backcross) |
| Hatfield and Schluter, 1999 | Benthic-limnetic | 6 populations (limnetic, limnetic backcross, F1 hybrid, F2 hybrid, benthic, benthic backcross) |
| Boughman et al., 2004 | Benthic-limnetic | 6 populations (3 benthic, 3 limnetic populations) |
| Hendry et al., 2011 | Lake-stream | 2 populations |
| Svanbäck and Schluter, 2012 | Benthic-limnetic | 2 populations |
| DeFaveri and Merilä, 2013 | Salinity levels | 6 populations (2 high salinity, 2 mid salinity, 2 low salinity) |
| Hudson et al, 2023 | Freshwater lineages | 2 populations (1 older, 1 newer) |
| Oke et al., 2016 | Lake-stream | 3 populations |
| Pilakouta et al., 2023 | Temperature | 12 populations (6 body of waters, 2 temperatures each) |

**Supplementary Table S2:** Population names, their abbreviation, and their sample sizes for trait study and fibrosis and infection study.

| Population name | Population abbreviation | Coordinates | Trait study sample size | Fibrosis and infection study sample size |
| --- | --- | --- | --- | --- |
| Amor Lake | AMO | 50°9' 57.96" N, 125°33' 14.4036" W | 21 | 150 |
| Anutz Lake | ANU | 50°18' 4.68" N, 126°55' 22.7856" W | 34 | 157 |
| Beaver Lake | BEA | 50°36' 2.52" N, 127°18' 54.0072" W | 31 | 43 |
| Black Lake | BLA | 48°46' 25.68" N, 125°5' 49.2" W | 24 | 162 |
| Blackwater Lake | BLK | 50°10' 18.4764" N, 125°35' 23.9856" W | 23 | 89 |
| Boot Lake | BOO | 50°3' 14.0472" N, 125°31' 51.5928" W | 45 | 86 |
| Brewster Lake | BRE | 50°6' 3.6072" N, 125°34' 58.8" W | 18 | 97 |
| Browns Bay Lake | BRB | 50°9' 32.0436" N, 125°24' 46.8072" W | 19 | 108 |
| Comida Lake | COM | 50°8' 58.2072" N, 125°31' 47.9964" W | 73 | 91 |
| Echo Lake | ECH | 49°59' 9.6" N, 125°24' 39.6108" W | 29 | 104 |
| Farewell Lake | FAR | 50°12' 0.7164" N, 125°35' 23.9856" W | 25 | 50 |
| Fredrick Lake | FRE | 48°51' 15.4836" N, 125°1' 33.6036" W | 14 | 64 |
| Gray Lake | GRY | 50°3' 25.5528" N, 125°35' 45.6036" W | 35 | 154 |
| Kidney Lake | KID | 50°36' 24.8364" N, 127°20' 2.4" W | 61 | 42 |
| Skating Pond Lake | L2B | 48°48' 52.9236" N, 125°7' 33.6" W | 27 | 1223 |
| Lawson Lake | LAW | 50°2' 9.5964" N, 125°34' 29.9892" W | 65 | 150 |
| Little Mud Lake | LMU | 50°12' 22.32" N, 125°33' 0.0108" W | 19 | 89 |
| Little Woss Lake | LWO | 50°10' 51.2436" N, 126°36' 43.1964" W | 58 | 154 |
| Merrill Lake | MER | 50°3' 41.0436" N, 125°33' 46.8108" W | 6 | 152 |
| Nimpkish Lake | NIM | 50°24' 37.44" N, 126°58' 29.9928" W | 13 | 107 |
| Ormond Lake | ORM | 50°10' 49.0872" N, 125°31' 33.6036" W | 4 | 90 |
| Pachena Lake | PCH | 48°50' 15.0072" N, 125°1' 44.3964" W | 13 | 52 |
| Roselle Lake | ROS | 50°31' 14.5164" N, 126°59' 20.3928" W | 82 | 101 |
| Rosseau Lake | RSS | 48°49' 49.4364" N, 124°59' 42.0108" W | 24 | 167 |
| Sarita Lake | SRA | 48°54' 33.4872" N, 124°53' 6.0072" W | 69 | 138 |
| Thiemer Lake | THE | 50°30' 37.8072" N, 126°58' 44.3856" W | 93 | 156 |
| Woss Lake | WOS | 50°7' 13.4364" N, 126°36' 46.7964" W | 37 | 65 |

**Supplementary Table S3:** Loadings of morphological traits on linear discriminant (LD) axes 1–14 derived from size-adjusted trait values, along with the proportion of trace explained by each axis. Trait loadings represent the contribution of each variable to the corresponding discriminant axis, with larger absolute values indicating stronger influence. The proportion of trace indicates the relative amount of among-population variation explained by each axis, with earlier axes accounting for the majority of variation and thus representing the primary dimensions of morphological divergence. Analysis of variance (ANOVA) results for LD axes derived from size-adjusted morphological traits. F-statistics and corresponding *p* values are reported for each LD axis, testing for differences among populations. All tests were conducted with 26 degrees of freedom. Significant differentiation among populations was observed along LD1 through LD10 (p < 0.05), including after Sequential Bonferroni correction (in bold) and for LD11 before Sequential Bonferroni correction (*) indicating that multiple axes of multivariate trait variation contribute to population divergence, while higher-order axes were not statistically significant.

| Traits | LD1 | LD2 | LD3 | LD4 | LD5 | LD6 | LD7 | LD8 | LD9 | LD10 | LD11 | LD12 | LD13 | LD14 |
| --- | --- | --- | --- | --- | --- | --- | --- | --- | --- | --- | --- | --- | --- | --- |
| Standard Length | 0.0100 | 1.1510 | 0.6242 | 0.1581 | -0.0550 | -0.2222 | -0.4944 | 0.2508 | 0.0007 | -0.0106 | -0.1895 | 0.0343 | 0.0620 | -0.0220 |
| Head Length | -0.5447 | -0.3896 | 0.5086 | 0.1914 | 0.2891 | -0.4205 | -0.0537 | -0.0780 | 1.3682 | 0.0661 | 0.6823 | -0.2584 | 0.8562 | -0.1057 |
| Snout Length | 0.1476 | 0.0770 | -0.8403 | -0.7209 | 0.0794 | -0.0525 | -0.6633 | -0.0462 | -1.2311 | -0.1488 | -0.7735 | -0.3648 | -0.2206 | -0.2121 |
| Eye Diameter | 0.2246 | 0.1879 | 0.4629 | -0.7268 | -0.7967 | 0.0796 | 0.6186 | -0.2227 | -0.3627 | -0.1415 | -0.0933 | 0.1797 | 0.1199 | -0.0458 |
| Caudal Peduncle Depth | -0.7552 | -0.3155 | -0.2307 | 0.4345 | -0.8024 | 0.0828 | -0.0805 | 0.7286 | -0.002 | 0.3816 | -0.3674 | 0.1867 | -0.0754 | 0.0384 |
| Body Depth | 1.3692 | 0.6614 | -0.5721 | 0.4086 | 0.0948 | 0.4797 | 0.5166 | 0.2365 | 0.1085 | -0.6326 | 0.1061 | -0.1419 | 0.1681 | -0.1372 |
| Gape Width | -0.0269 | 0.1419 | -0.6167 | 0.0908 | -0.0169 | -0.4231 | -0.3608 | -0.7097 | 0.3600 | 0.3182 | 0.1083 | 0.6394 | -0.0321 | 0.2597 |
| Opercular 4-bar KT | 0.0744 | 0.1876 | -0.2200 | -0.0989 | -0.0412 | -0.2973 | -0.0566 | 0.0831 | -0.1402 | 0.3841 | 0.6848 | -0.3331 | -0.3276 | 0.0944 |
| Length Gill Raker 1 | -0.0157 | -0.2550 | -0.0045 | -0.2497 | 0.1672 | -0.6500 | 0.2936 | 0.2359 | -0.0933 | 0.0847 | 0.2126 | 0.8439 | -0.5828 | -1.8475 |
| Length Gill Raker 2 | 0.3046 | -0.165 | 0.1203 | -0.2195 | 0.2407 | 0.4191 | 0.3075 | 0.4145 | 0.1552 | -0.3258 | 0.0184 | -0.2414 | 0.3755 | 2.0847 |
| Gill Raker Density | -0.3214 | 0.1746 | -0.0286 | 0.1643 | -0.0619 | -0.1163 | 0.0886 | -0.2288 | 0.2941 | 0.7719 | -0.1224 | -0.2607 | -0.5941 | 0.1165 |
| Fin Aspect Ratio | 0.1948 | -0.1693 | -0.1693 | 0.1341 | 0.1238 | -0.7375 | 0.5356 | 0.2654 | -0.0392 | 0.3117 | -0.4305 | -0.3173 | -0.0908 | 0.1038 |
| Armor Plate N | 0.6513 | -0.7232 | 0.5261 | 0.4448 | -0.7139 | -0.3719 | -0.5682 | -0.2259 | -0.508 | -0.0894 | -0.1694 | 0.0183 | 0.0263 | -0.0532 |
| Gill Raker N | 0.1310 | 0.0631 | -0.1488 | -0.5397 | -0.1859 | 0.1148 | -0.0336 | 0.2654 | -0.0508 | 0.5365 | -0.5236 | -0.0431 | -0.4527 | 0.0087 |
| Proportion of trace | 0.3406 | 0.1794 | 0.1476 | 0.1034 | 0.0681 | 0.0407 | 0.0344 | 0.0259 | 0.0177 | 0.0144 | 0.0116 | 0.009 | 0.0053 | 0.0020 |
| *F* value, *p* value | **51.46,**<br>**<0.0001** | **27.11,**<br>**<0.0001** | **22.30,**<br>**<0.0001** | **15.63,**<br>**<0.0001** | **10.28,**<br>**<0.0001** | **6.15,**<br>**<0.0001** | **5.191,**<br>**<0.0001** | **3.91,**<br>**<0.0001** | **2.671,**<br>**0.0001** | **2.179,**<br>**0.0008** | 1.752,<br>0.0135 * | 1.356,<br>0.116 | 0.807,<br>0.739 | 0.304,<br>1 |

**Supplementary Table S4:** Confusion matrix from the LDA of size-adjusted morphological traits. Rows indicate the true (actual) population assignment of each individual, and columns indicate the population predicted by the LDA. Values along the diagonal (in green) represent correctly classified individuals, whereas off-diagonal values represent misclassified individuals (in gray).

|  |  |  |  |  |  |  |  |  |  |  |  |  |  |  |  |  |  |  |  |  |  |  |  |  |  |  |  |
| --- | --- | --- | --- | --- | --- | --- | --- | --- | --- | --- | --- | --- | --- | --- | --- | --- | --- | --- | --- | --- | --- | --- | --- | --- | --- | --- | --- |
| WOS | 0 | 0 | 0 | 0 | 0 | 0 | 0 | 0 | 0 | 1 | 0 | 1 | 0 | 0 | 1 | 0 | 0 | 0 | 0 | 4 | 0 | 0 | 0 | 0 | 2 | 2 | 11 |

**Supplementary Table S5:**
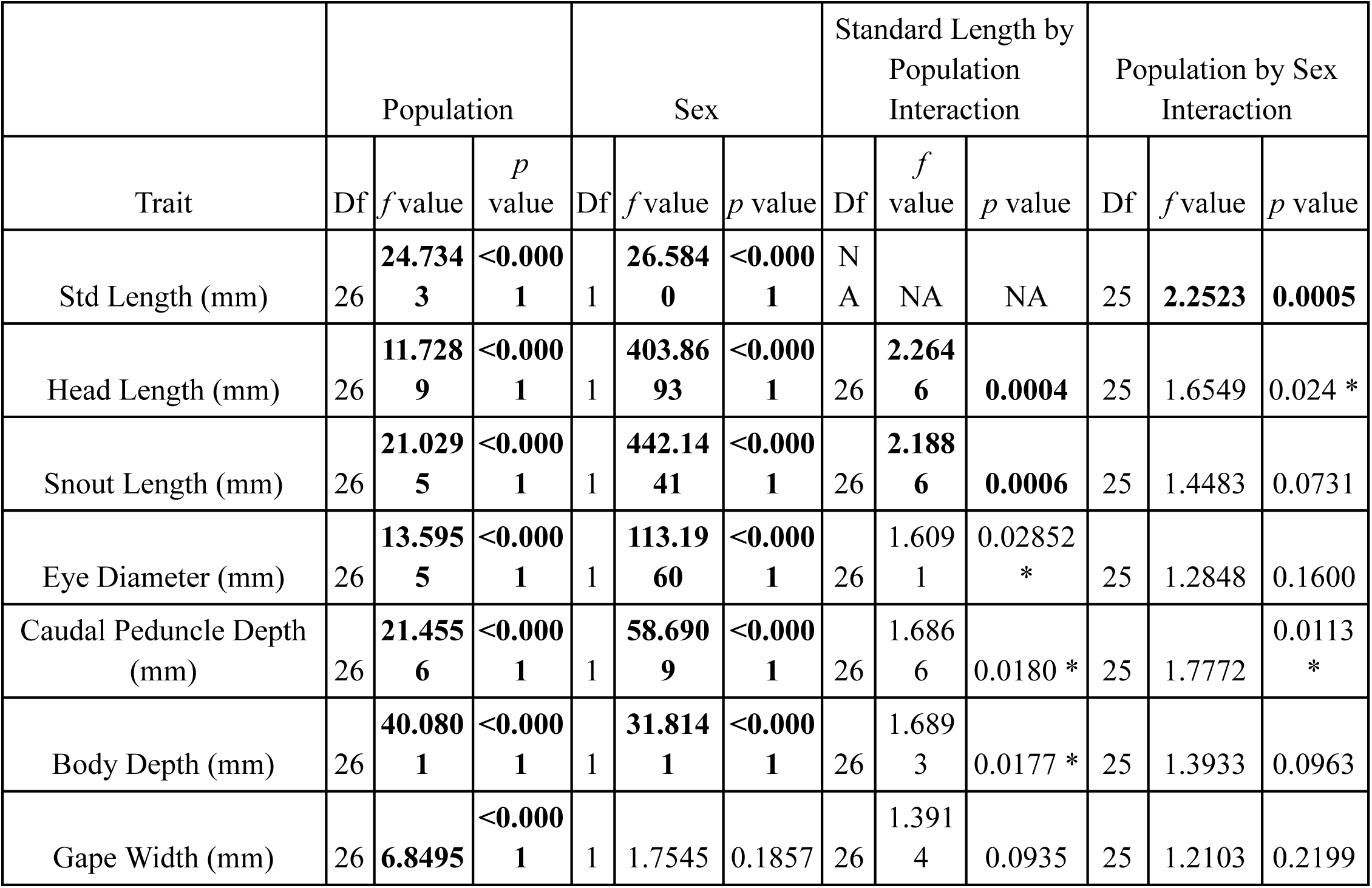
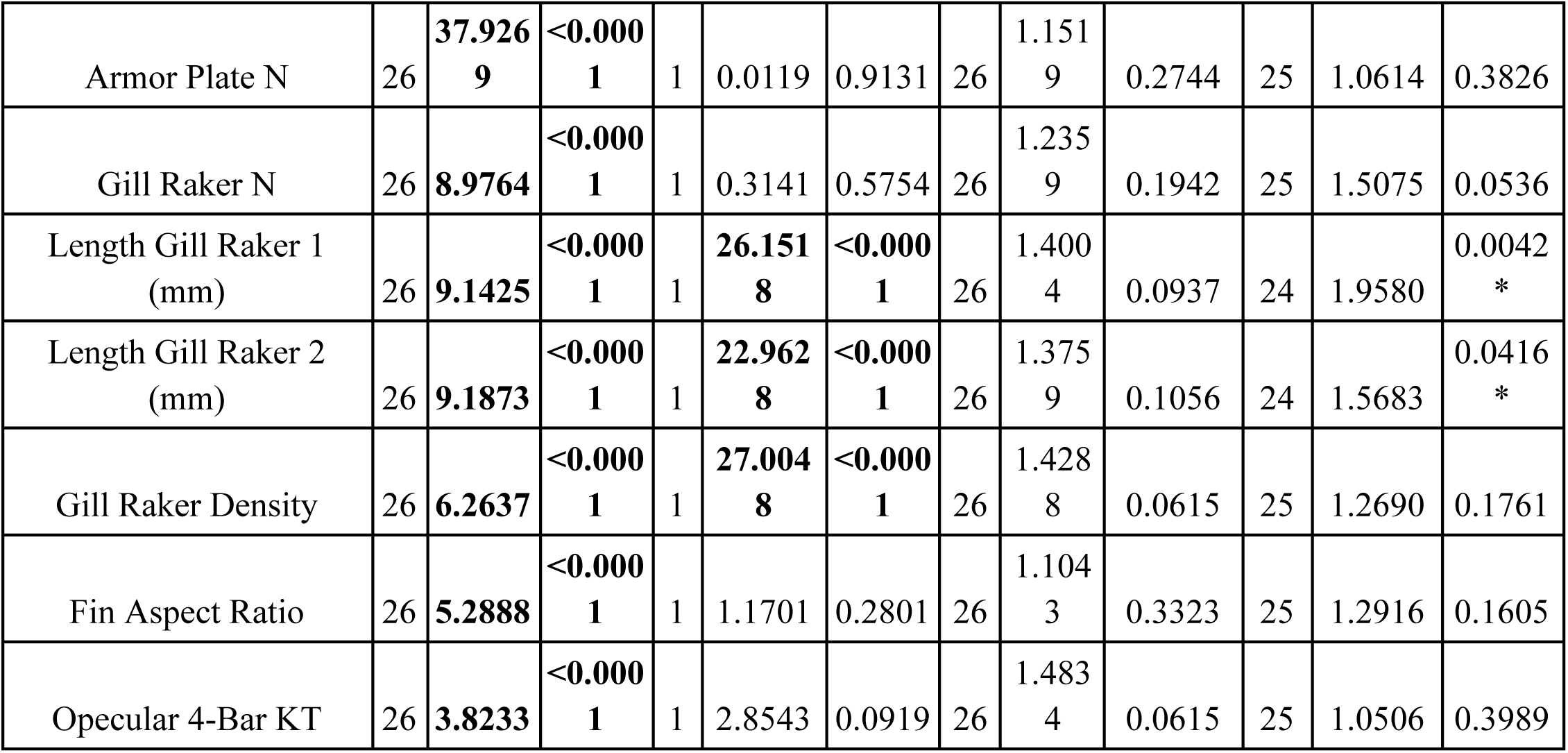
Results of linear models testing the effects of population, sex, and body size (standard length) on morphological traits. Traits were size-adjusted where appropriate, except for standard length, armor plate number, gill raker number, and fin aspect ratio. For each trait, *f* statistics and corresponding *p* values are reported for the main effects of population and sex, as well as for interaction between standard length by population and sex by population. Significant effects indicate variation in trait values attributable to differences among populations, between sexes, or differences in allometric scaling across populations and sexes. Bolded are significant after Sequential Bonferroni (*P* < 0.05). An asterisk (*) within the *p* value indicates significance at *p* < 0.05.

**Supplementary Table S6:**
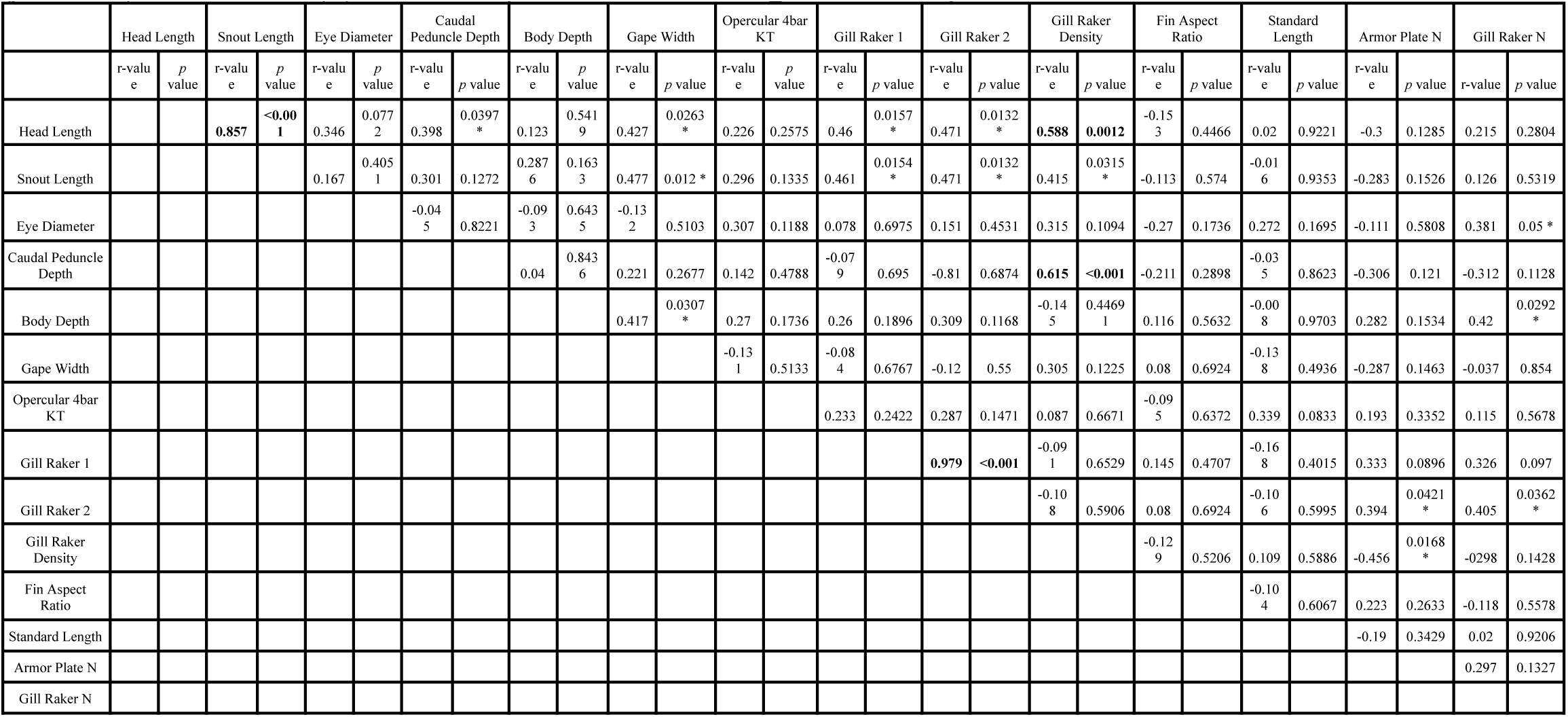
Correlations between morphological traits. Those that are bolded are significant after Sequential Bonferroni (*p* < 0.05). An asterisk (*) within the *p* value indicates significance at *p* < 0.05.

**Supplementary Table S7:**
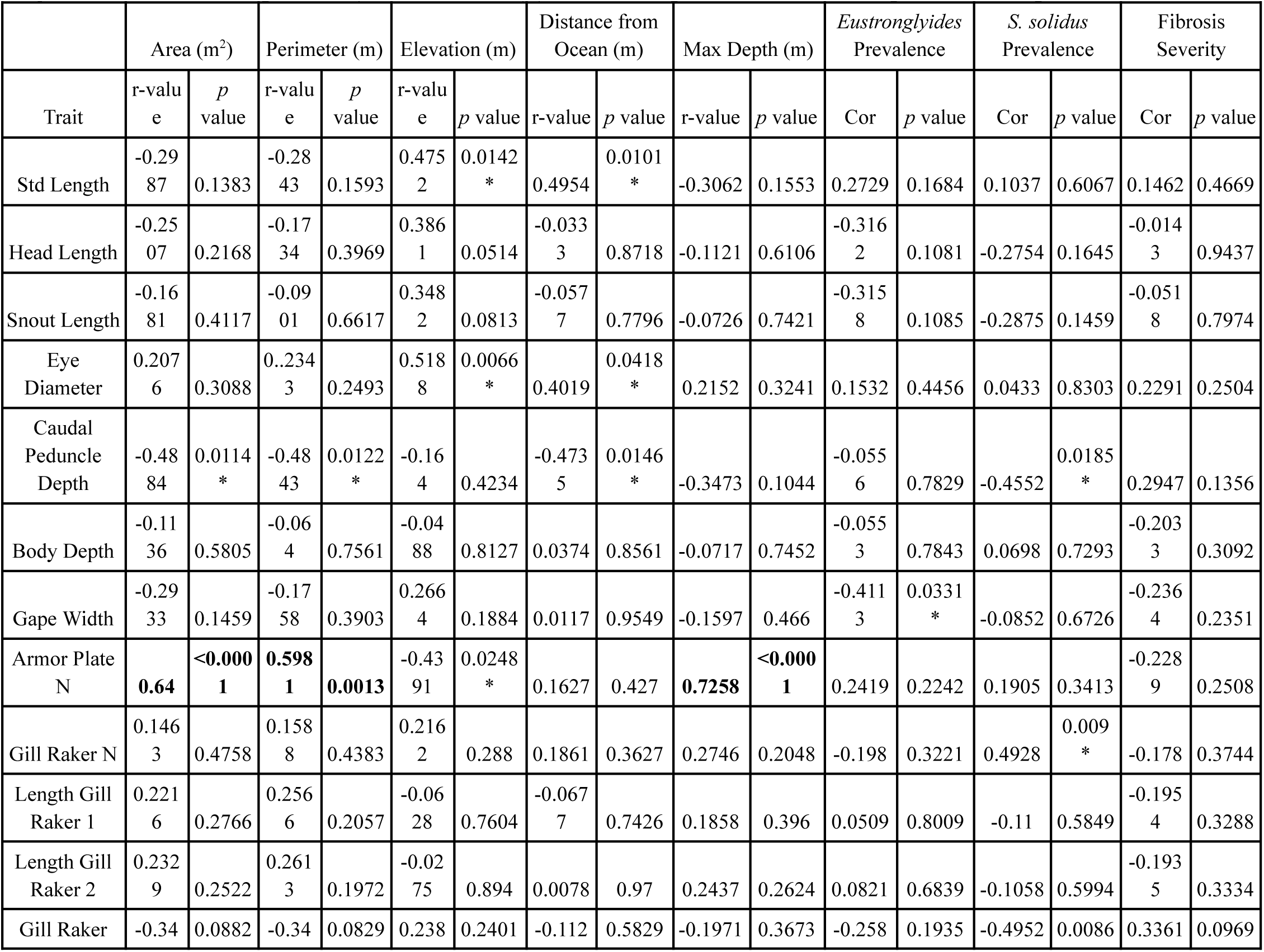
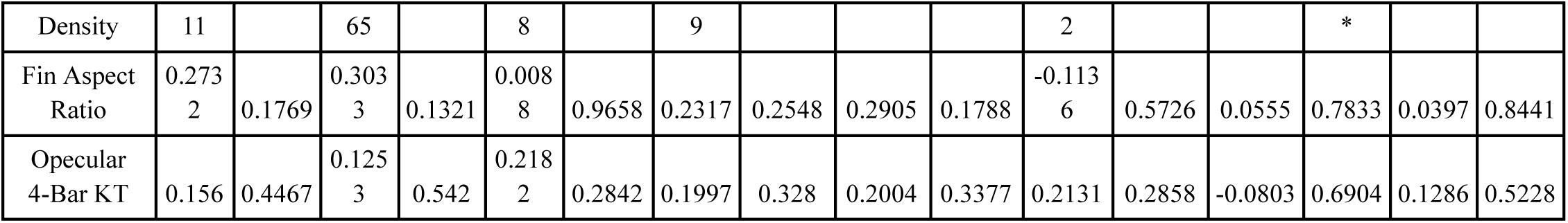
Correlations between lake environmental variables (log transformed) and parasite prevalence (including *Eustrongylides*, *S. solidus*, and fibrosis) with population mean morphological traits. Pearson correlation coefficients (r) are reported for each pairwise comparison, with corresponding *p* values indicating statistical significance. Those that are bolded are significant after Sequential Bonferroni (*p* < 0.05). An asterisk (*) within the *p* value indicates significance at *p* < 0.05.

**Supplementary Table S8:** Results of linear regression analyses examining relationships between morphological traits and environmental covariates across stickleback populations. For each significant model, the table reports the morphological trait, predictor category (landscape and climate or water column), environmental predictor variable, standardized regression coefficient (Beta), standard error (SE), *t*-statistic, and *p* value.

| Trait | Category | Predictor | Beta | SE | t stat | *p* val |
| --- | --- | --- | --- | --- | --- | --- |
| Armor Plate N | Landscape & Climate | Max Depth (Est.) [m] | 0.993 | 0.141 | 7.02 | < 0.001 |
| Armor Plate N | Landscape & Climate | Degree Days <18°C (Spring) | 0.23 | 0.107 | 2.16 | 0.043 |
| Armor Plate N | Landscape & Climate | Lake Perimeter [m] | 1.532 | 0.156 | 9.83 | < 0.001 |
| Body Depth | Landscape & Climate | Tertiary Sp. Cover [%] | 0.28 | 0.106 | 2.64 | 0.015 |
| Body Depth | Landscape & Climate | Precipitation (Jun) [mm] | 0.253 | 0.105 | 2.41 | 0.024 |
| Body Depth | Landscape & Climate | Moisture Index (Dec) [mm] | 0.225 | 0.108 | 2.08 | 0.048 |
| CP Depth | Water Column (Sonde) | Dissolved O2 SD [%] | 0.047 | 0.016 | 3 | 0.007 |
| CP Depth | Landscape & Climate | Frost-Free Days (Summer) | 0.047 | 0.017 | 2.67 | 0.014 |
| CP Depth | Landscape & Climate | Dist. to Paved Road [m] | -0.065 | 0.018 | -3.59 | 0.002 |
| CP Depth | Landscape & Climate | Rel. Humidity (Spring) [%] | 0.049 | 0.02 | 2.43 | 0.023 |
| CP Depth | Landscape & Climate | Paved Road Density [km/km<sup>2</sup>] | 0.05 | 0.019 | 2.61 | 0.016 |
| CP Depth | Water Column (Sonde) | fDOM SD [RFU] | 0.059 | 0.019 | 3.12 | 0.005 |
| Eye Diameter | Landscape & Climate | Rel. Humidity (Feb) [%] | 0.061 | 0.015 | 4.13 | < 0.001 |
| Eye Diameter | Landscape & Climate | Dist. to Nearest Lake [m] | -0.046 | 0.019 | -2.47 | 0.021 |
| Eye Diameter | Landscape & Climate | Degree Days <18°C (Sep) | 0.059 | 0.015 | 3.84 | < 0.001 |
| Eye Diameter | Landscape & Climate | mean DD 0 02 | 0.05 | 0.016 | 3.18 | 0.004 |
| Eye Diameter | Landscape & Climate | Rel. Humidity (Nov) [%] | 0.062 | 0.015 | 4.2 | < 0.001 |
| Eye Diameter | Landscape & Climate | Rel. Humidity (Dec) [%] | 0.058 | 0.015 | 3.75 | 0.001 |
| Eye Diameter | Landscape & Climate | Crown Closure [%] | 0.038 | 0.017 | 2.24 | 0.036 |
| Eye Diameter | Landscape & Climate | Degree Days <18°C (Feb) | 0.043 | 0.017 | 2.53 | 0.019 |
| Eye Diameter | Landscape & Climate | Rel. Humidity (Jan) [%] | 0.062 | 0.015 | 4.23 | < 0.001 |
| Gill Raker Density | Landscape & Climate | Hargreaves Ref. Evap. (Oct) [mm] | -0.009 | 0.004 | -2.08 | 0.048 |
| Gill Raker Density | Water Column (Sonde) | Water Temp SD [°C] | 0.012 | 0.003 | 3.37 | 0.003 |
| Gill Raker Density | Landscape & Climate | Dist. to Paved Road [m] | -0.009 | 0.004 | -2.34 | 0.029 |
| Gill Raker Density | Landscape & Climate | Hargreaves Ref. Evap. (Autumn) [mm] | -0.011 | 0.004 | -2.63 | 0.015 |
| Gill Raker Density | Landscape & Climate | Degree Days <18°C (Sep) | -0.01 | 0.004 | -2.63 | 0.015 |
| Gill Raker Density | Landscape & Climate | Rel. Humidity (Oct) [%] | 0.012 | 0.004 | 3.22 | 0.004 |
| Gill Raker Density | Landscape & Climate | Snow Precip. (Spring) [mm] | -0.011 | 0.004 | -2.97 | 0.007 |
| Gill Raker 1 length | Landscape & Climate | Dominant Sp. Cover [%] | -0.047 | 0.018 | -2.66 | 0.014 |
| Gill Raker 1 length | Landscape & Climate | Dist. to Ancient Forest [m] | 0.044 | 0.016 | 2.68 | 0.013 |
| Gill Raker 1 length | Landscape & Climate | Vertical Complexity | -0.052 | 0.016 | -3.3 | 0.003 |
| Gill Raker 1 length | Water Column (Sonde) | pH | 0.05 | 0.014 | 3.58 | 0.002 |
| Gill Raker 1 length | Landscape & Climate | Tertiary Sp. Cover [%] | 0.054 | 0.017 | 3.07 | 0.005 |
| Gill Raker 1 length | Landscape & Climate | Degree Days <18°C (Spring) | 0.06 | 0.015 | 4.03 | < 0.001 |
| Gill Raker 1 length | Landscape & Climate | Forest Basal Area [m<sup>2</sup>/ha] | 0.071 | 0.013 | 5.27 | < 0.001 |
| Gill Raker 2 length | Landscape & Climate | Forest Basal Area [m<sup>2</sup>/ha] | 0.074 | 0.012 | 6.01 | < 0.001 |
| Gill Raker 2 length | Landscape & Climate | Dist. to Ancient Forest [m] | 0.052 | 0.017 | 3.03 | 0.006 |
| Gill Raker 2 length | Landscape & Climate | Vertical Complexity | -0.051 | 0.017 | -3.06 | 0.006 |
| Gill Raker 2 length | Water Column (Sonde) | pH | 0.037 | 0.014 | 2.57 | 0.018 |
| Gill Raker 2 length | Landscape & Climate | Tertiary Sp. Cover [%] | 0.042 | 0.018 | 2.34 | 0.029 |
| Gill Raker 2 length | Landscape & Climate | Dominant Sp. Cover [%] | -0.051 | 0.017 | -2.94 | 0.007 |
| Gill Raker 2 length | Landscape & Climate | Degree Days <18°C (Spring) | 0.061 | 0.015 | 4.14 | < 0.001 |
| Gill Raker N | Landscape & Climate | Elevation [m] | 0.393 | 0.188 | 2.09 | 0.048 |
| Gill Raker N | Water Column (Sonde) | fDOM [RFU] | -0.62 | 0.166 | -3.73 | 0.001 |
| Head Length | Landscape & Climate | Rel. Humidity (Feb) [%] | 0.175 | 0.049 | 3.55 | 0.002 |
| Head Length | Landscape & Climate | Crown Closure [%] | 0.127 | 0.054 | 2.34 | 0.028 |
| Head Length | Landscape & Climate | Rel. Humidity (Jan) [%] | 0.173 | 0.05 | 3.47 | 0.002 |
| Head Length | Landscape & Climate | Hargreaves Ref. Evap. (Oct) [mm] | -0.16 | 0.052 | -3.09 | 0.005 |
| Head Length | Landscape & Climate | Rel. Humidity (Winter) [%] | 0.169 | 0.05 | 3.36 | 0.003 |
| Opercular 4bar KT | Landscape & Climate | Moisture Index (Aug) [mm] | -0.169 | 0.055 | -3.07 | 0.005 |
| Opercular 4bar KT | Landscape & Climate | Forest Basal Area [m²/ha] | 0.213 | 0.055 | 3.83 | < 0.001 |
| Opercular 4bar KT | Landscape & Climate | Degree Days <18°C (Spring) | 0.184 | 0.06 | 3.07 | 0.005 |
| Opercular 4bar KT | Landscape & Climate | Hargreaves Deficit (Aug) [mm] | 0.171 | 0.055 | 3.11 | 0.005 |
| Opercular 4bar KT | Landscape & Climate | Dist. to Nearest Lake [m] | -0.168 | 0.06 | -2.8 | 0.01 |
| Opercular 4bar KT | Landscape & Climate | Canopy Height [m] | 0.174 | 0.053 | 3.26 | 0.003 |
| Opercular 4bar KT | Landscape & Climate | Stand Height [m] | 0.174 | 0.053 | 3.26 | 0.003 |
| Opercular 4bar KT | Landscape & Climate | Rel. Humidity (Nov) [%] | 0.193 | 0.057 | 3.4 | 0.002 |
| Opercular 4bar KT | Landscape & Climate | Max Temp (Dec) [°C] | -0.15 | 0.057 | -2.61 | 0.016 |
| Snout Length | Landscape & Climate | Crown Closure [%] | 0.091 | 0.039 | 2.34 | 0.028 |
| Snout Length | Landscape & Climate | Degree Days <18°C (Jun) | 0.111 | 0.037 | 2.99 | 0.007 |
| Snout Length | Landscape & Climate | Stand Height [m] | 0.097 | 0.038 | 2.54 | 0.018 |
| Snout Length | Landscape & Climate | Forest Basal Area [m²/ha] | 0.105 | 0.037 | 2.82 | 0.01 |
| Snout Length | Landscape & Climate | Canopy Height [m] | 0.097 | 0.038 | 2.54 | 0.018 |
| Snout Length | Landscape & Climate | Degree Days >5°C (Jun) | 0.101 | 0.038 | 2.63 | 0.016 |
| Snout Length | Landscape & Climate | Mean Temp (Jun) [°C] | 0.1 | 0.038 | 2.61 | 0.016 |
| Snout Length | Landscape & Climate | Degree Days <18°C (Jun) | -0.101 | 0.038 | -2.63 | 0.016 |
| Standard Length | Landscape & Climate | Snow Precip. (May) [mm] | -2.553 | 0.727 | -3.51 | 0.002 |
| Standard Length | Landscape & Climate | Forest Harvest Year | 2.656 | 0.671 | 3.96 | < 0.001 |
| Standard Length | Landscape & Climate | Precipitation (May) [mm] | -2.812 | 0.723 | -3.89 | < 0.001 |
| Standard Length | Landscape & Climate | Rel. Humidity (Feb) [%] | 1.835 | 0.818 | 2.24 | 0.035 |
| Standard Length | Landscape & Climate | Degree Days <18°C (Autumn) | 1.947 | 0.77 | 2.53 | 0.019 |
| Standard Length | Landscape & Climate | Rel. Humidity (Winter) [%] | 1.956 | 0.803 | 2.44 | 0.023 |
| Gape Width | None | (No significant predictors found) | - | - | - | - |
| Fin Aspect Ratio | None | (No significant predictors found) | - | - | - | - |

**Supplementary Table S9:** Results of the distance-based redundancy analysis (dbRDA) examining whether variation in zooplankton community composition explains multivariate morphological variation among common-garden reared stickleback populations. (A) Variance explained by the dbRDA model, including raw and adjusted *R*² values. (B) Global permutation test evaluating the significance of the overall constrained model. (C) Marginal permutation tests evaluating the independent contribution of each zooplankton community principal component (PC1 and PC2) to morphological variation.

| Metric | Value |
| --- | --- |
| R-Squared (Raw) | 0.0827 |
| Adjusted R-Squared | 0.0029 |
| Model Significance<br>($p$-value) | 0.3890 |

B: dbRDA global permutation
| Effect | Df | Variance | F | Pr(>F) |
| --- | --- | --- | --- | --- |
| Model | 2 | 1.1572 | 1.0362 | 0.3980 |
| Residual | 23 | 12.8428 | - | - |

C: dbRDA marginal axis
| Axis | Df | Variance | F | Pr(>F) |
| --- | --- | --- | --- | --- |
| Zoop_Community_PC1 | 1 | 0.5568 | 0.9971 | 0.4400 |
| Zoop_Community_PC2 | 1 | 0.5734 | 1.0268 | 0.4150 |
| Residual | 23 | 12.8428 | - | - |

## Appendix Table

**Appendix Table 1:**
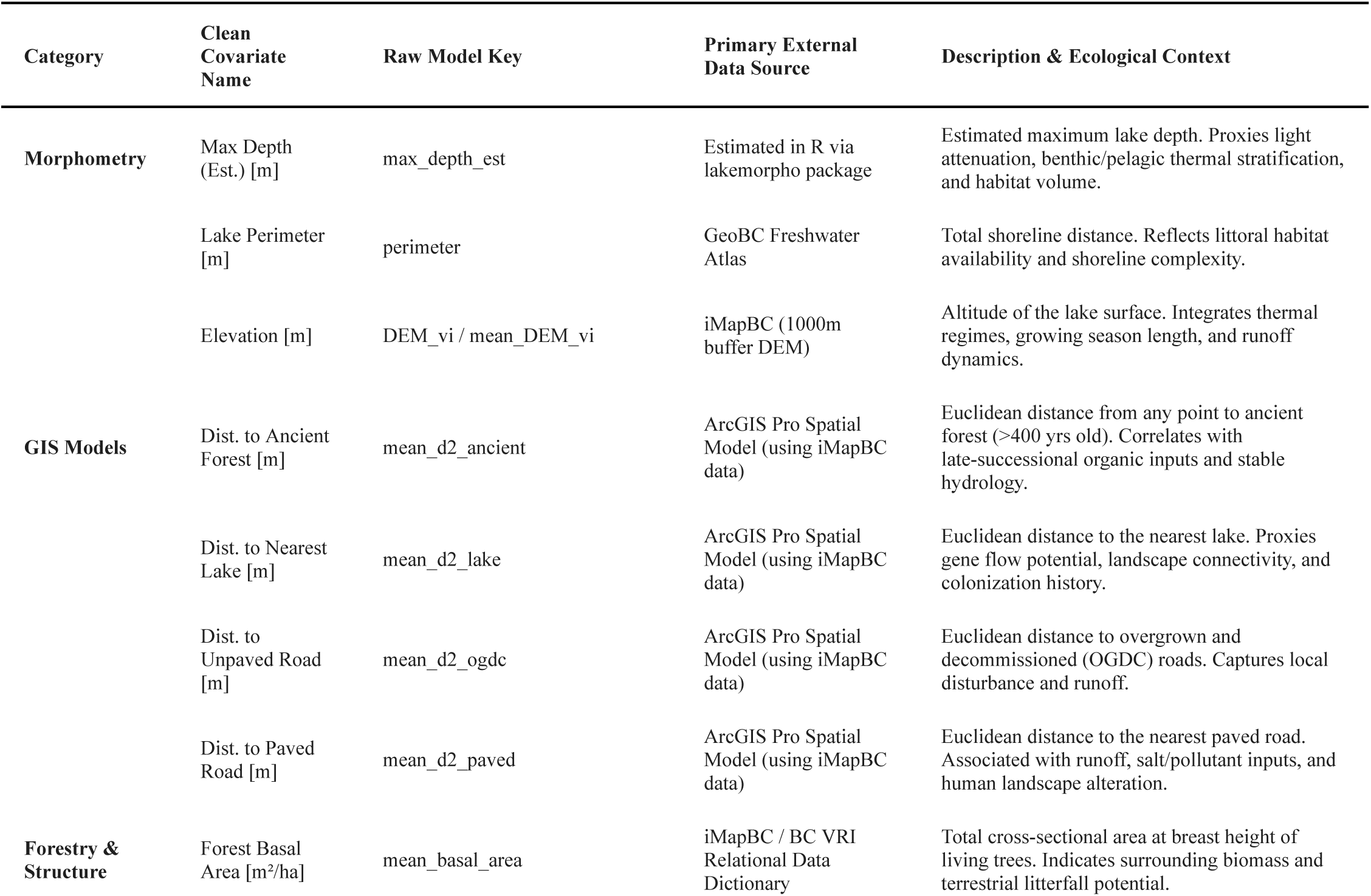
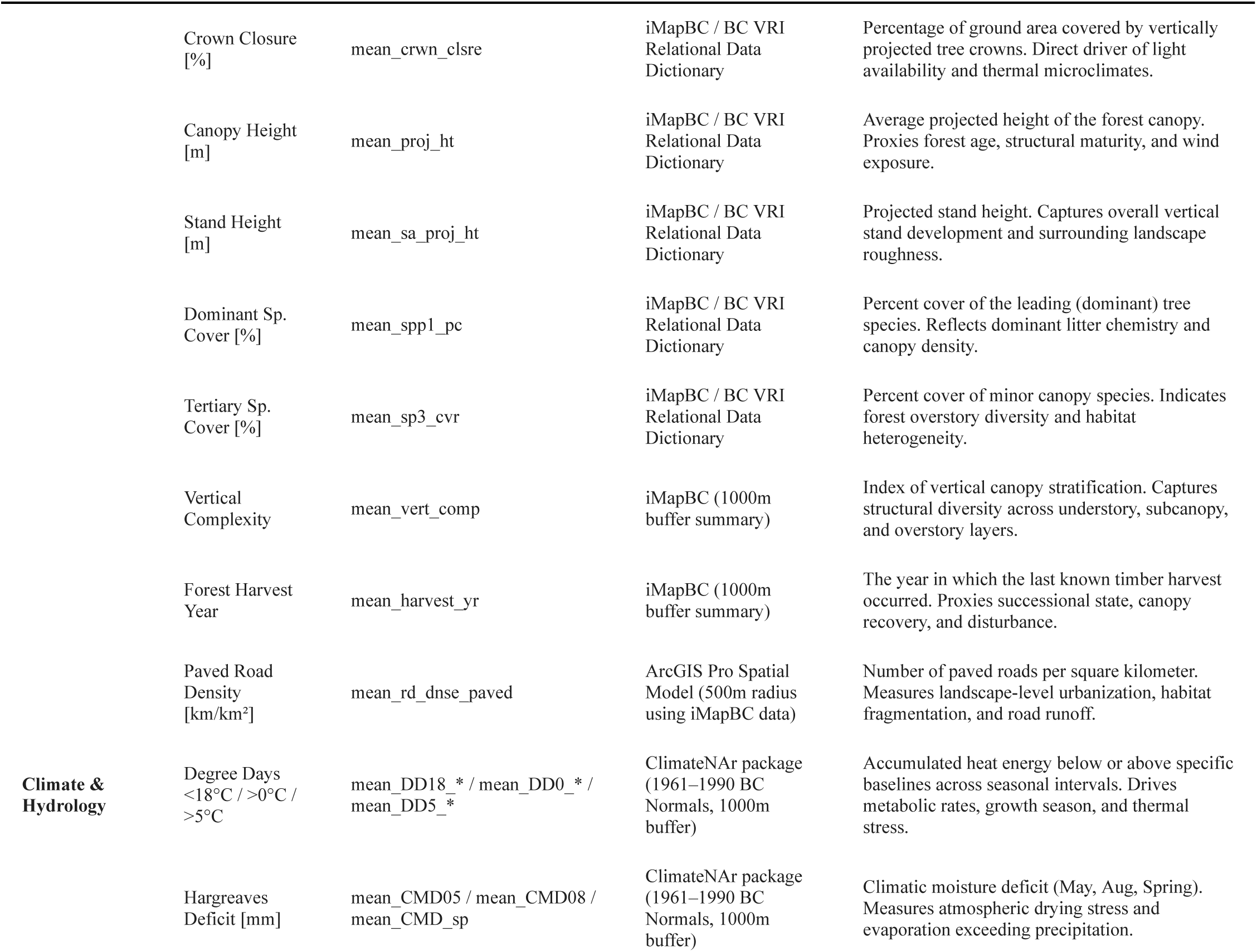
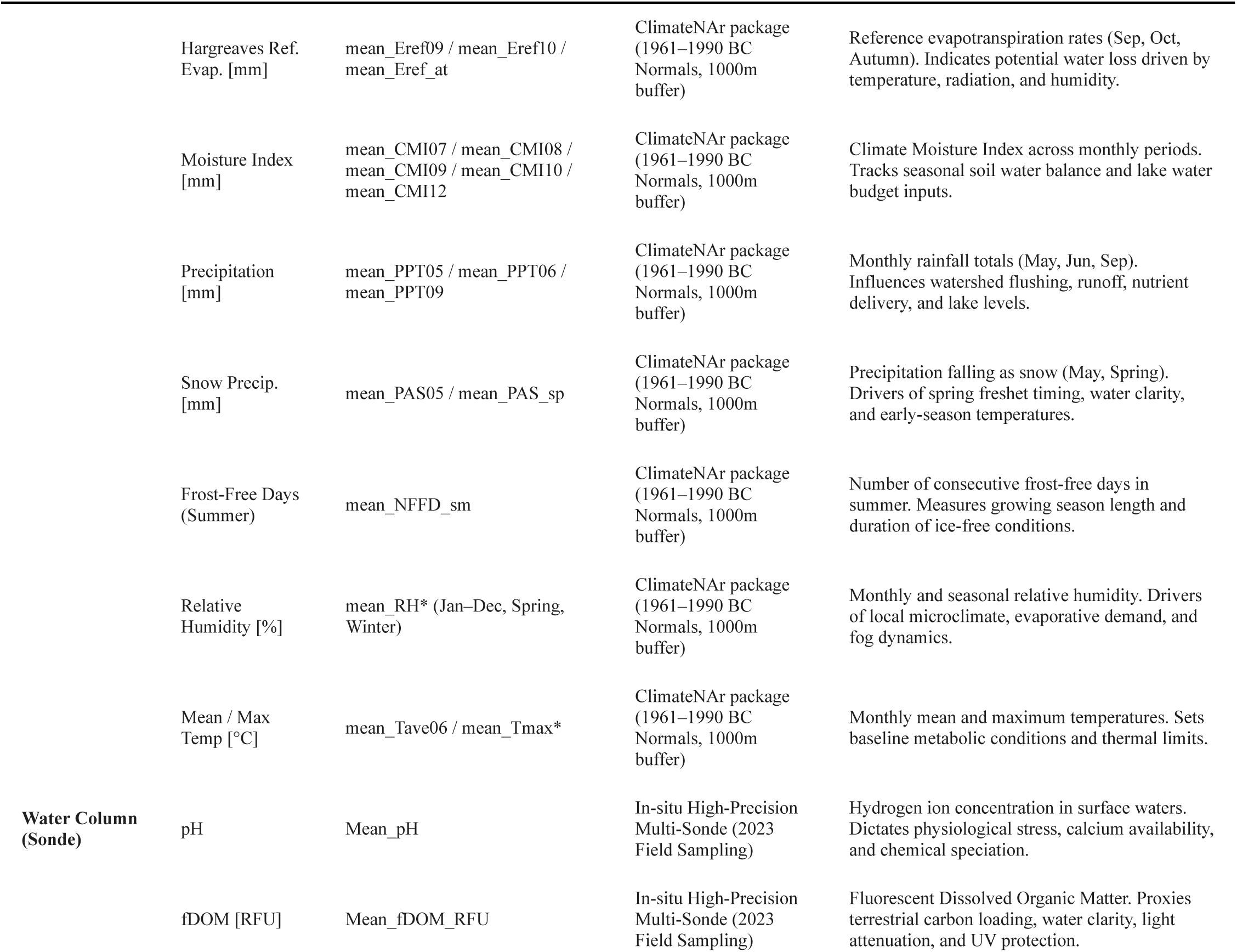
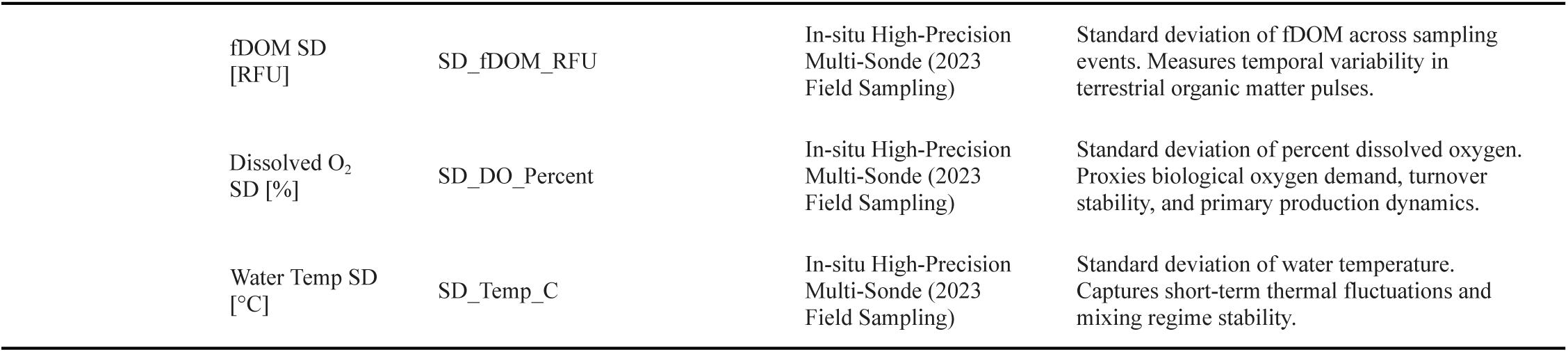
Environmental covariates category, predictors raw and clean name, primary external data source, and description and ecological context of variables.

## Supplements

### Supplement 1: Methods for Environmental Covariates

To evaluate drivers of morphology, we paired site-level fish morphometrics with spatially explicit environmental data, including landscape, climate, water column, and zooplankton community data. We summarized multi-depth water column profiles into lake-level means and standard deviations using the *group by* and *summarise* functions from the *dplyr* package (Wickham et al., 2023).

We applied several specialized functions from the vegan package to account for site-specific variations in the zooplankton community (Oksanen et al., 2022). First, we used the *decostand* function to perform a Hellinger transformation on raw zooplankton taxon counts per lake, stabilizing site-to-site variation in rare species abundances. We then performed a Principal Component Analysis (PCA) via the *rda* function. Using the *scores* function, we extracted the site-specific PCA coordinates along the first two principal component axes (*PC1* and *PC2*) to use as continuous site-level predictors reflecting landscape-level gradients in the zooplankton community.

To prevent missing data from forcing us to exclude valuable biological observations, we addressed gaps in our predictive environmental matrix using a two-tiered preprocessing protocol. First, we imputed missing values in continuous environmental features using a baseline mean replacement approach. This step substituted empty cells with the calculated average value of that specific environmental feature across all other sampled lakes, preserving our overall sample size without shifting any variable’s global mean. This strategy affected a minor portion of our data, totaling only 144 missing numeric cells out of 14,006 available data points (1.03% of the numeric matrix). Second, we handled missing entries within categorical features by dynamically assigning unobserved values to an explicit category level labeled ‘Missing’ via the *fct_na_value_to_level* function from the *forcats* package (Wickham, 2023).

Following data imputation, we applied two structural filtering criteria to clean the predictor pool before running our models. We discarded any environmental variable exhibiting zero variance, as features without variation offer no statistical leverage to explain differences in fish morphology. Furthermore, we excluded complex categorical predictors containing more than three unique factor levels. This specific restriction safeguarded our analysis against downstream computational bias, preventing machine learning architectures like Random Forests from becoming artificially skewed toward selecting these high-cardinality categories over truly meaningful physical or biological drivers.

Because our cumulative environmental data pool contained a large, highly correlated set of predictors (*n = 276*) relative to our individual biological measurements (*n = 13*), we implemented a non-parametric variable selection workflow. For each distinct fish morphological trait (e.g., standard length, fin width, eye diameter), we used the *ranger* function within the *ranger* package to fit independent Random Forest models configured with 1,000 decision trees (Wright and Ziegler, 2017). We set the split-variable parameter value (the *mtry* argument within the *ranger* function) to one-third of the total predictor pool to balance tree strength and independence (Breiman, 2001). We isolated individual feature importance using non-parametric permutation variable importance scores, truncated negative importance metrics to zero, and transformed individual scores into relative percentage contributions. We selected environmental features for subsequent confirmatory modeling if their mean relative importance surpassed a dynamic threshold defined as *100/N_vars_* (where *N_vars_* represents the total number of input variables evaluated), retaining a maximum of the top ten predictors per trait.

We evaluated the environmental predictors selected during our Random Forest screening against their respective stickleback morphological traits using independent linear regression models via the *lm* function from the *stats* package (R Core Team, 2026). Prior to model fitting, we *z*-score standardized all continuous predictors using the *scale* function in the base package of R, converting our raw slopes into standardized effect sizes (*b*). We handled categorical factors using standard dummy coding (treatment contrasts), which evaluated all group differences relative to an alphabetical baseline category. For every candidate model, we calculated Cook’s Distance metrics across all individual fish observations via the *cooks.distance* function in the *stats* package (R Core Team, 2026). If an observation exceeded our operational leverage threshold of *4/N* (where *N* represents the model’s specific sample size), which serves as a standard benchmark for identifying highly influential data points (Bollen and Jackman, 1985), we removed it to prevent individual outliers from biasing our results. Next, we refitted our linear models using these filtered datasets to compute final standardized slopes (*b*), standard errors (*SE*), test statistics (*t*), and parametric *p*-values, using a significance threshold of *a* = 0.05. Finally, we generated analysis of variance (ANOVA) tables using the *anova* function (*stats*) to evaluate the overarching significance of our multi-level categorical factors.

To complement our linear models and verify the global influence of zooplankton communities on stickleback morphology, we performed a distance-based Redundancy Analysis (dbRDA) using the *dbrda* function within the *vegan* package (Oksanen et al., 2022). We first constructed a morphological response matrix containing all scaled stickleback traits. Then, using the *vegdist* function, also within the vegan package, we calculated a multivariate Euclidean distance matrix directly from these traits to quantify phenotypic dissimilarity among individuals. Finally, we modeled this resulting distance matrix as a joint function of our extracted zooplankton community axes (*PC1* + *PC2*). We calculated both the total (*R^2^*) and adjusted variation (*R^2^_adj_*) explained using the *RsquareAdj* function. To determine if global morphological patterns were statistically meaningful, we tested the overall significance of this multi-trait space using 999 random permutations via the *anova.cca* function, which simultaneously allowed us to isolate the unique, independent effect of each zooplankton axis.

